# Pathway and network analysis of more than 2,500 whole cancer genomes

**DOI:** 10.1101/385294

**Authors:** Matthew A. Reyna, David Haan, Marta Paczkowska, Lieven P.C. Verbeke, Miguel Vazquez, Abdullah Kahraman, Sergio Pulido-Tamayo, Jonathan Barenboim, Lina Wadi, Priyanka Dhingra, Raunak Shrestha, Gad Getz, Michael S. Lawrence, Jakob Skou Pedersen, Mark A. Rubin, David A. Wheeler, Søren Brunak, Jose MG Izarzugaza, Ekta Khurana, Kathleen Marchal, Christian von Mering, S. Cenk Sahinalp, Alfonso Valencia, Jüri Reimand, Joshua M. Stuart, Benjamin J. Raphael, on behalf of the PCAWG Drivers and Functional Interpretation Group and the ICGC/TCGA Pan-Cancer Analysis of Whole Genome Network

## Abstract

The catalog of cancer driver mutations in protein-coding genes has greatly expanded in the past decade. However, non-coding cancer driver mutations are less well-characterized and only a handful of recurrent non-coding mutations, most notably *TERT* promoter mutations, have been reported. Motivated by the success of pathway and network analyses in prioritizing rare mutations in protein-coding genes, we performed multi-faceted pathway and network analyses of non-coding mutations across 2,583 whole cancer genomes from 27 tumor types compiled by the ICGC/TCGA PCAWG project. While few non-coding genomic elements were recurrently mutated in this cohort, we identified 93 genes harboring non-coding mutations that cluster into several modules of interacting proteins. Among these are promoter mutations associated with reduced mRNA expression in *TP53, TLE4*, and *TCF4*. We found that biological processes had variable proportions of coding and non-coding mutations, with chromatin remodeling and proliferation pathways altered primarily by coding mutations, while developmental pathways, including Wnt and Notch, altered by both coding and non-coding mutations. RNA splicing was primarily targeted by non-coding mutations in this cohort, with samples containing non-coding mutations exhibiting similar gene expression signatures as coding mutations in well-known RNA splicing factors. These analyses contribute a new repertoire of possible cancer genes and mechanisms that are altered by non-coding mutations and offer insights into additional cancer vulnerabilities that can be investigated for potential therapeutic treatments.

## Introduction

Over the past decade, cancer genome sequencing efforts such as The Cancer Genome Atlas (TCGA) have identified millions of somatic genetic aberrations; however, the annotation and interpretation of these aberrations remains a major challenge^1^. Specifically, while some aberrations occur frequently in specific cancer types, there is a “long tail” of rare aberrations that are difficult to distinguish from random passenger aberrations in modestly sized patient cohorts^2,3^. In many cancers, a significant proportion of patients do not have known coding driver mutations^4^, suggesting that additional driver mutations remain undiscovered. To date, the vast majority of known driver mutations affect protein-coding regions; only a few non-coding driver mutations, most notably mutations in the *TERT* promoter^5–7^, have been identified. Recent studies from the Pan-Cancer Analysis of Whole Genomes (PCAWG) project of the International Cancer Genome Consortium (ICGC) reveal few recurrent non-coding drivers in analyses of individual genes and regulatory regions^7^.

Cancer driver mutations unlock oncogenic properties of cells by altering the activity of hallmark pathways^8^. Accordingly, cancer genes are known to cluster in small number of cellular pathways and interacting subnetworks^3,9^. Previously, pathway and network analysis has proven useful for implicating infrequently mutated genes as cancer genes based on their pathway membership and physical/regulatory interactions with recurrently mutated genes^10–14^. However, the interactions between coding and non-coding driver mutations have not been systematically explored.

We performed pathway and network analysis of coding and non-coding somatic mutations from 2,583 tumors from 27 tumor types compiled by the Pan-Cancer Analysis of Whole Genomes (PCAWG) project of the International Cancer Genome Consortium (ICGC)^15^, the largest collection of uniformly processed cancer genomes to date. We derive a consensus set of 93 high-confidence pathway-implicated driver genes with non-coding variants (PID-N) and a consensus set of 87 pathway-implicated driver genes with coding variants (PID-C) using seven pathway and network analysis methods. Both sets of PID genes, particularly the PID-N set, contain rarely mutated genes that were not identified by individual recurrence tests but interact with other well-known cancer genes. In total, 121 novel PID-N and PID-C genes are revealed as promising candidates, expanding the landscape of driver mutations in cancer.

Furthermore, we examined the contribution of coding and non-coding mutations in altering biological processes, finding that while chromatin remodeling and some well-known signaling and proliferation pathways are altered primarily by coding mutations, other important cancer pathways, including developmental pathways such as Wnt and Notch pathways, are altered by both coding and non-coding mutations in PID genes. Intriguingly, we find many non-coding mutations in PID-N genes with roles in RNA splicing, and samples with these non-coding mutations exhibit similar gene expression signatures as samples with well-known coding mutations in RNA splicing factors. Our analysis demonstrates that somatic non-coding mutations in untranslated and cis-regulatory regions constitute a complementary set of genetic perturbations with respect to coding mutations, affect several biological pathways and molecular interaction networks, and should be further investigated for their role in the onset and progression of cancer.

## Results

### The long tail of coding and non-coding cancer mutations highlights opportunities for pathway and network analysis

We analyzed the genes targeted by single nucleotide variants (SNVs) and short insertions and deletions (indels) identified by whole genome sequencing in the 2,583 ICGC PCAWG tumor samples from 27 tumor types. Our pathway and network analyses focused on a subset of 2,252 tumors that excluded melanomas and lymphomas due to their atypical distributions of mutations in regulatory regions^16^. We analyzed the pan-cancer driver *p*-values of single protein-coding and non-coding elements predicted by the PCAWG consensus driver analysis^7^ including exons, promoters, untranslated regions (5’ UTR and 3’ UTR), and enhancers. This PCAWG consensus driver analysis integrates *p*-values from 16 driver discovery methods, resulting in consensus driver *p*-values for coding and non-coding elements. Among protein-coding driver *p*-values of the pan-cancer cohort, 75 genes were highly significant (FDR < 0.1; **Supplemental Figure S1**) and an additional 7 genes were observed at near-significant levels (0.1 ≤ FDR < 0.25). These numbers are consistent with previous reports of a “long tail” of driver genes with few highly-mutated genes and many genes with infrequent mutations across cancer types^2,17^. Non-coding mutations exhibit a similar long-tail distribution with even fewer significant genes (8 genes at FDR < 0.1 and 2 genes at 0.1 ≤ FDR < 0.25). No single gene has both significant or near-significant coding and non-coding driver *p*-values (FDR < 0.25), suggesting that non-coding mutations target a complementary set of genes as coding mutations.

Earlier studies have demonstrated that proteins harboring coding driver mutations interact with each other in molecular pathways and networks significantly more frequently than expected by chance^2,3,9–11,13^. We observed significant numbers of interactions between both coding and/or non-coding elements with more mutations than expected by chance, suggesting that pathway and network methods may be able to identify rare driver events that are not prioritized by single-element analyses (**Supplemental Figure S2; *Coding and non-coding mutations cluster on networks* in Supplement**).

### Consensus pathway and network analysis reveals possible non-coding driver mutations

We performed a comprehensive pathway and network analysis of cancer drivers using the results of the single-element driver discovery study of the PCAWG project^7^ as input. Our methods leveraged prior pathway and network knowledge to amplify the results of this single-element analysis. We performed a consensus analysis from seven distinct methods (ActivePathways [Paczkowska, Barenboim *et al*., in submission], CanIsoNet [Kahraman *et al*., in preparation], Hierarchical HotNet^18^, a hypergeometric analysis [Vazquez], an induced subnetwork analysis [Reyna and Raphael, in preparation], NBDI^19^, and SSA-ME^20^) that utilized information from molecular pathways or protein interaction networks (**Figure 1, Methods**). Each method nominated genes, and consensus sets of genes with possible coding and non-coding driver mutations were defined as the genes found by at least four of the seven methods (**Supplemental Tables S1-S4**). All methods were calibrated on randomized data (***Individual pathway and network algorithms* in Supplement**).

**Figure 1:**
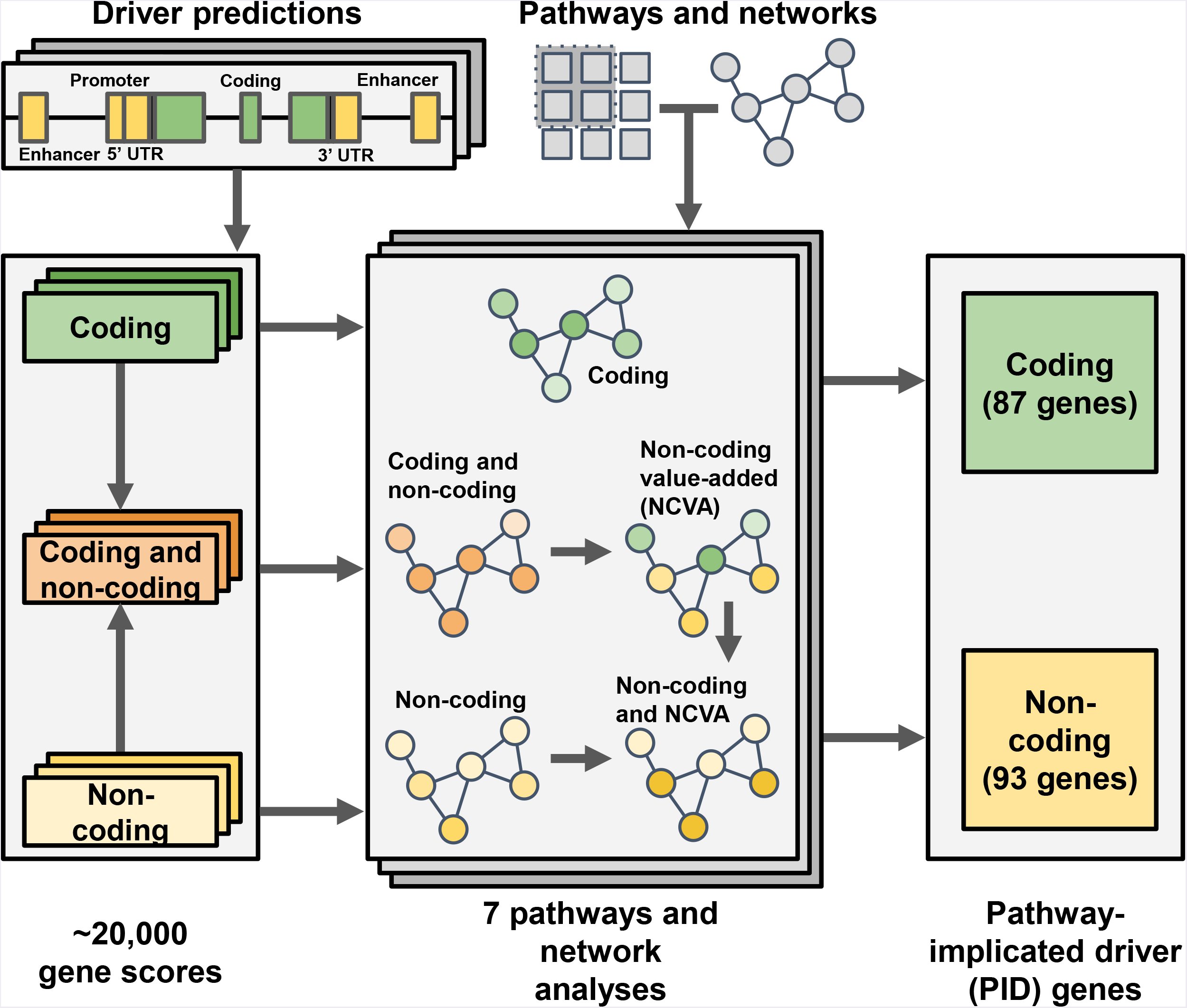
Overview of the pathway and network analysis approach. Coding, non-coding, and combined gene scores were derived for each gene by aggregating driver *p*-values from the PCAWG driver predictions in individual elements, including annotated coding and non-coding elements (promoter, 5’ UTR, 3’ UTR, and enhancer). These gene scores were input to five network analysis algorithms, which utilize multiple protein-protein interaction networks, and to two pathway analysis algorithms, which utilize multiple pathway/gene-set databases. We defined a non-coding value-added (NCVA) procedure to determine genes whose non-coding scores contribute significantly to the results of the combined coding and non-coding analysis, where NCVA results for a method augment its results on non-coding data. We defined a consensus procedure to combine significant pathways and networks identified by these seven algorithms. The 87 pathway-implicated driver genes with coding variants (PID-C) are the set of genes reported by a majority (≥ 4/7) of methods on coding data. The 93 pathway-implicated driver genes with non-coding variants (PID-N) are the set of genes reported by a majority of methods on non-coding data or in their NCVA results.

When using non-coding mutations alone, a consensus of pathway and network analysis results on non-coding data identified 62 genes. In contrast, and as one might expect, the coding analysis resulted in substantially more genes, producing a set of 87 pathway-implicated driver genes with coding variants (PID-C). To increase the sensitivity for detecting contributions provided by non-coding mutations, we devised a “non-coding value-added” (NCVA) procedure (**Figure 1, Supplemental Figure S3; *Non-coding value-added (NCVA) procedure* in Methods**). Our NCVA procedure asks if the coding mutations enhance the discovery of potential non-coding driver genes beyond what is found with only the non-coding mutations. This procedure identified an additional set of 31 genes that, when merged with the 62 genes found with non-coding mutations alone, resulted in a set of 93 pathway-implicated driver genes with non-coding variants (PID-N) (**Supplemental Figure S4, *Consensus results* in Methods**). PID-N genes appear as a robust and biologically relevant set, unbiased by any particular mutational process reflecting a particular carcinogen or DNA damage processes (**Supplemental Figure S5, *Mutational signatures* in Methods**).

The 87 PID-C genes (**Supplementary Table 1, Supplemental Figure S6A**) include 68 previously identified cancer genes as catalogued by the COSMIC Cancer Gene Census (CGC) database (v83, 699 genes from Tier 1 and Tier 2)^21^ (2.98 genes expected; Fisher’s exact test *p* = 3.57 × 10^-83^; **Figures 2A and 2C, Supplemental Figure S7A**). The PID-C genes have significantly higher coding gene scores than non-PID-C genes (rank sum test *p* = 1.72 × 10^-58^; median rank 48 of PID-C genes), and each of the 87 PID-C genes improves the score of its network neighborhood (19.7 genes expected; *p* < 10^-6^; **Supplemental Table S5**). This network neighborhood analysis shows that PID-C genes are not implicated solely by their network neighbors^14^ but themselves contribute significantly to their discovery by pathway and network methods. The 87 PID-C genes also include 31 genes that are not statistically significant (FDR > 0.1) in the PCAWG single-element driver analysis; **Figures 2A and 2C; Supplemental Figures S8A and S9**), illustrating that the network neighborhoods can nominate genes with infrequent mutations, i.e., those in the “long tail”, as possible driver genes. Interestingly, 13 of these 31 genes with FDR > 0.1 are also known drivers according to the CGC database (3.0 genes expected; Fisher’s exact test *p* = 2.1 × 10^-14^). Thus, the consensus pathway and network analysis recovers many known protein-coding driver mutations and identifies additional possible drivers that are infrequently mutated and thus remain below the statistical significance threshold of gene-specific driver analyses.

**Figure 2:**
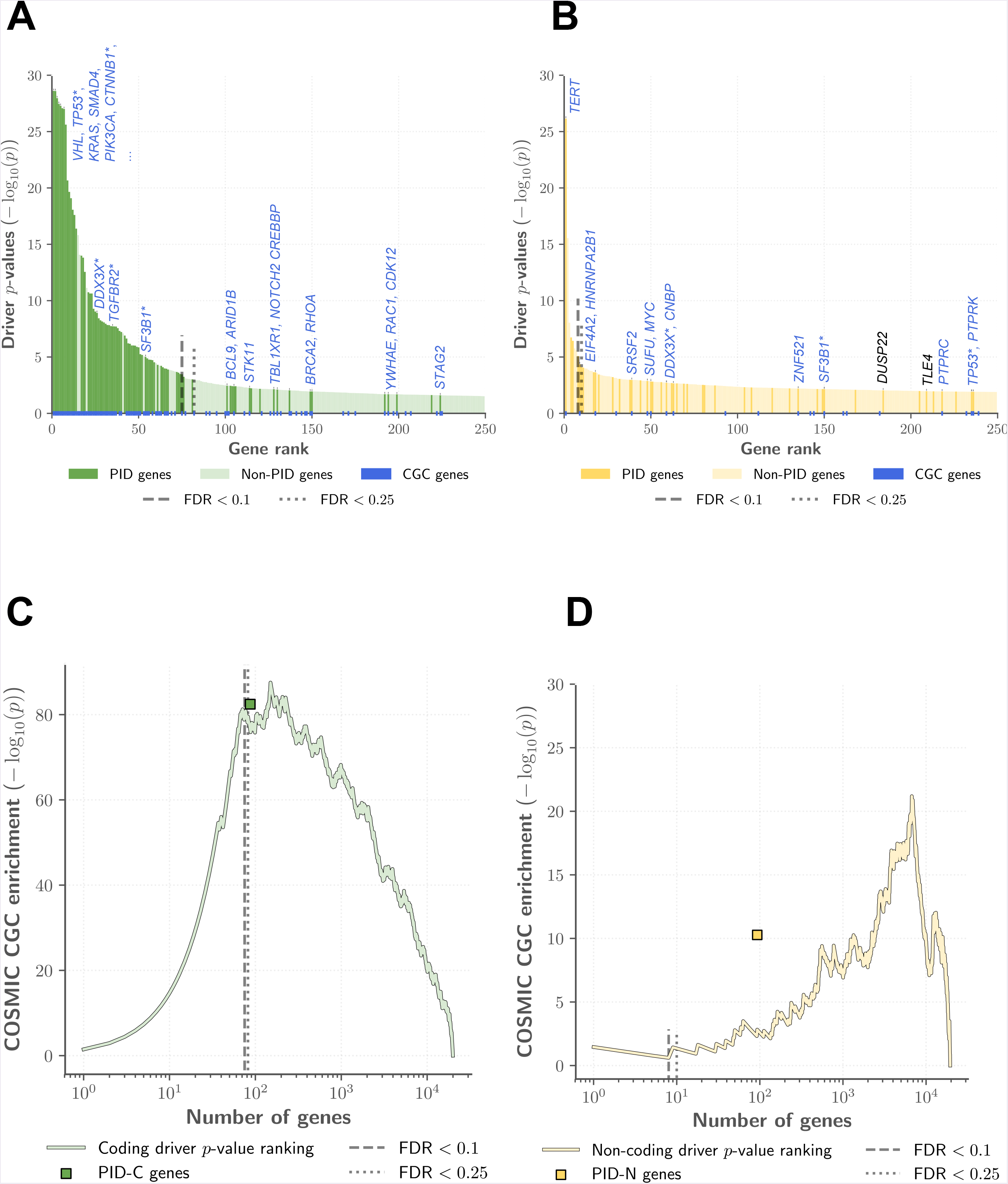
**(A) Pathway and network methods identify significant coding driver mutations**. Driver *p*-values on protein-coding elements for the 250 genes with most significant coding driver *p*-values; dashed and dotted lines indicate FDR = 0.1 and 0.25, respectively. Dark green bars are PID-C genes, while light green bars are non-PID-C genes. Blue squares below the x-axis indicate genes from the COSMIC Cancer Gene (CGC) Census. In total, 31 of 87 PID-C genes have coding driver *p*-values with FDR > 0.1 and would not be reported as drivers using single-gene tests with the typical FDR = 0.1 threshold^4,7,14^. Several PID-C genes are labeled, including all COSMIC CGC genes with coding FDR > 0.1. Genes that are both PID-C and PID-N genes are indicated with asterisks. Note that 3 PID-C genes are not among the 250 most significantly mutated genes shown in the figure. **(B) Pathway and network methods identify rare non-coding driver mutations**. Driver *p*-values on non-coding elements (promoter, 5’ UTR, and 3’ UTR of gene) for 250 genes with most significant non-coding driver *p*-values; dashed and dotted lines indicate FDR = 0.1 and 0.25, respectively. Dark yellow bars are PID-N genes, while light yellow bars are non PID-N genes. Blue squares below the x-axis indicate genes from the COSMIC CGC. In total, 3 (*TERT, HES1*, *TOB1*) of 93 PID-N genes have non-coding driver *p*-values with FDR ≤ 0.1, while 90 have FDR > 0.1, and thus would generally not be reported as drivers using single-gene tests. Several PID-N genes are labeled, including PID-N genes with significant *in cis* gene expression changes (**see Fig. 3**) and all PID-N genes with non-coding FDR > 0.25. Genes that are both PID-C and PID-N genes are indicated with asterisks. Note that 48 PID-N genes are not among the 250 most significantly mutated genes shown in figure. **(C)**. Statistical significance of overlap between top ranked genes according to coding driver *p*-values and PID-C genes with COSMIC Cancer Gene Census (CGC) genes. Overlap *p*-values are compute with Fisher’s exact test and driver FDR thresholds of 0.1 and 0.25 are highlighted. Green square indicates significance of overlap between PID-C genes and CGC genes. **(D)** Statistical significance of overlap of genes ranked by driver *p*-values on non-coding (promoter, 5’ UTR, 3’ UTR) elements and COSMIC CGC genes. Driver FDR thresholds of 0.1 and 0.25 are highlighted. Yellow square indicates significance of overlap between PID-N genes and CGC genes. Note the different scaling of y-axis compared to Fig. 2C.

The 93 PID-N genes (**Supplementary Table 2, Supplemental Figure S6B**) include 19 previously identified cancer genes according to the COSMIC Cancer Gene Census (CGC) database (3.2 genes expected; Fisher’s exact test *p* = 5.3 × 10^-11^; **Figures 2B and 2D; Supplemental Figures S7B and S7C**). Excluding the eight genes with individually significant non-coding elements from the PCAWG consensus drivers analysis^7^, 19 genes are both PID-N genes and CGC genes (3.1 genes expected; Fisher’s exact test *p* = 5.3 × 10^-11^), suggesting that non-coding mutations may alter genes with recurrent coding or structural variants in some samples. The PID-N genes have significantly higher non-coding gene scores than non-PID-N genes (rank sum test *p* = 1.47 × 10^-58^; median rank 165 of PID-N genes), and 92/93 PID-N (except for *HIST1H2BO*) genes improve the scores of their network neighborhoods (28.5 genes expected; *p* < 10^-6^; **Supplemental Table S6**). This network neighborhood analysis shows that PID-N genes are not implicated solely by their network neighbors^14^. The vast majority of PID-N genes (90 out of the 93, including the 19 CGC genes) are distinct from the PCAWG single-element driver analysis (**Figure 2B, Supplemental Figures S8B and S9**), with only three genes in common: *TERT, HES1, and TOB1*. Of these three, only *TERT* is recognized as a known driver according to the CGC database. Moreover, the 93 PID-N genes are more strongly enriched (Fisher’s exact test *p* = 5.3 × 10^-11^) for COSMIC CGC genes than the 93 genes with the smallest non-coding driver *p*-values of promoters, 5’ UTRs, or 3’ UTRs (Fisher’s exact test *p* = 4.8 × 10^-3^). Thus, our consensus procedure of the pathway and network analyses appreciably augments the PCAWG set of non-coding driver candidates.

Taken together, the PID-C and PID-N results identified an additional 121 genes over what was found in the element-focused PCAWG driver analysis, including 90 new possible non-coding drivers (***Consensus Results* in Methods**). In total, non-coding mutations in PID-N genes cover an additional 151 samples (9.1% of samples) than PID-C genes. In addition, the overwhelming majority of the PID-N genes were distinct from PID-C genes (88 out of 93; **Supplemental Figure S4**). While this suggests that coding and non-coding driver mutations have largely distinct gene targets, we show below that both types of mutations affect distinct sets of cancer genes underlying many of the same hallmark cancer processes.

### Impact of non-coding mutations on gene expression

As most PID-N genes have little support from previous studies to corroborate their roles in tumorigenesis, we sought to evaluate the biological relevance of the PID-N genes by testing whether non-coding mutations in a PID-N genes were associated with expression changes in that gene. Such *in cis* expression effects may be a result of the mutation located in transcription factor binding sites or other types of regulatory sites. We found that 5 PID-N genes (FDR < 0.3) showed statistically significant *in cis* correlations out of the 90 that could be tested using RNA-Seq data (**Figure 3; Supplemental Figure S10; Supplemental Tables S8-10, S12-14**). In contrast, 34 out of 87 PID-C genes with statistically significant or near statistically significant *in cis* expression changes (FDR < 0.3) (**Supplemental Tables S7, S11**).

**Figure 3:**
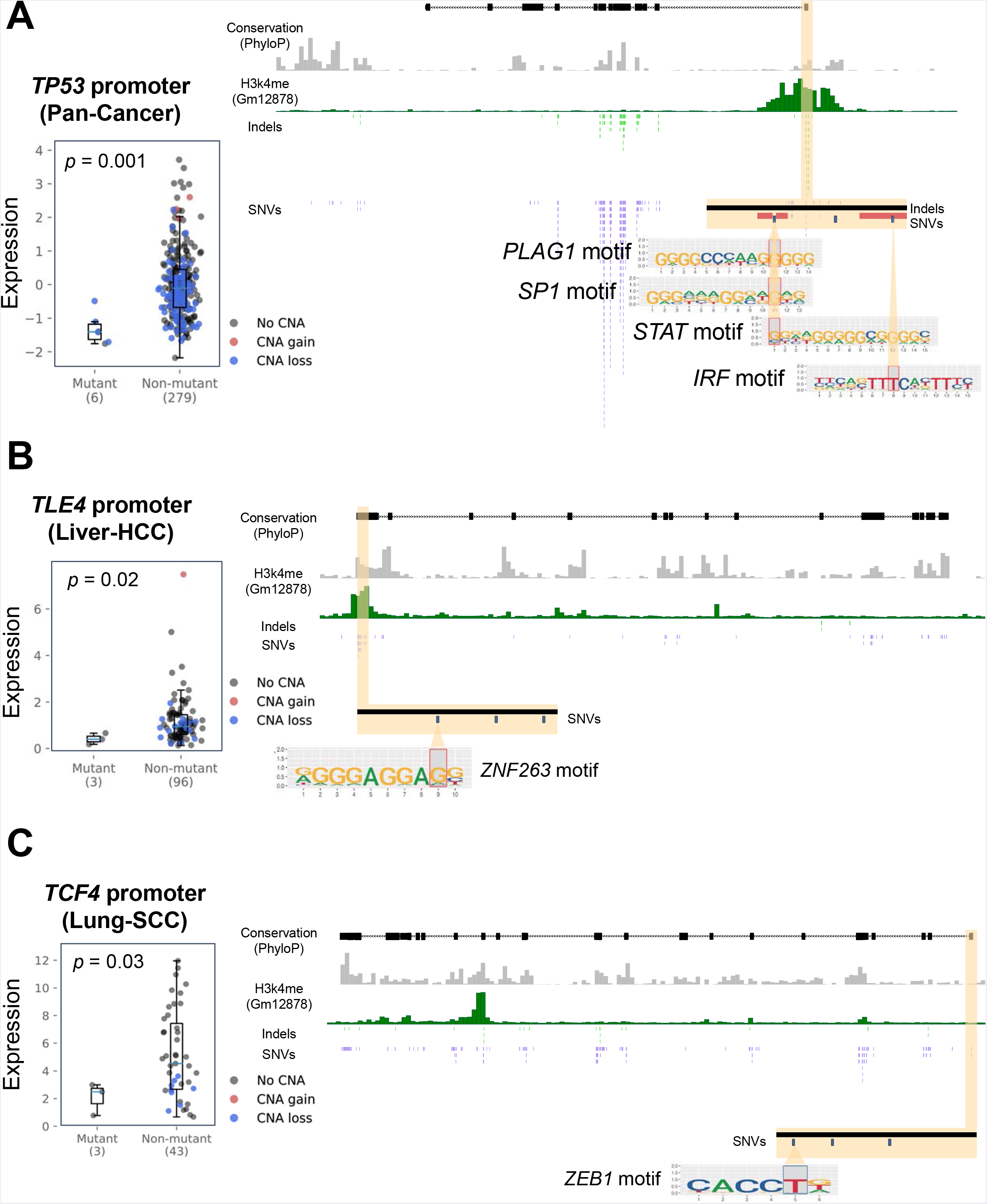
Gene expression changes are correlated with mutations in PID-N genes. **(A) *TP53* promoter**. *TP53* coding and non-coding genomic loci with zoomed-in view of *TP53* promoter region. *TP53* promoter mutations (six mutations in Biliary-AdenoCA, ColoRect-AdenoCA, Kidney-ChRCC, Lung-SCC, Ovary-AdenoCA, and Panc-AdenoCA cancer types) correlate significantly (Wilcoxon rank-sum test *p* = 0.001, FDR = 0.087) with reduced *TP53* gene expression. Samples with copy number gains and losses in the *TP53* promoter region are annotated in red and blue, respectively. Two of the six *TP53* promoter mutations overlap with transcription factor binding sites (with one mutation matching 3 motifs). **(B) *TLE4* promoter**. *TLE4* coding and non-coding genomic loci with zoomed-in view of *TLE4* promoter region. *TLE4* promoter mutations in Liver-HCC samples (three mutations) correlate (Wilcoxon rank-sum test *p* = 0.02, FDR = 0.2) with lower *TLE4* gene expression. Samples with copy number gains and losses annotated in red and blue, respectively. One of the three *TLE4* promoter mutations has a transcription factor binding site for *ZNF263*. **(C) *TCF4* promoter**. *TCF4* coding and non-coding genomic loci with zoomed-in view of *TCF4* promoter region. *TCF4* promoter mutations in Lung-SCC samples (three mutations) correlate (Wilcoxon rank-sum test *p* = 0.03, FDR = 0.27) with lower *TCF4* gene expression. Samples with copy number gains and losses annotated in red and blue, respectively. One of the the three *TCF4* promoter mutations has a transcription factor binding site for *ZEB1*.

Unsurprisingly, the most significant association between mutation and expression for PID-N genes is the correlation between *TERT* promoter mutations and increased expression, which we find in 11 Thy-AdenoCA tumors (Wilcoxon rank-sum test *p* = 1.3 × 10^-10^, FDR = 3.2 × 10^-9^), 11 CNS-Oligo tumors (Wilcoxon rank-sum test *p* = 6.8 × 10^-3^, FDR = 9.7 × 10^-2^), and 22 CNS-GBM tumors (Wilcoxon rank-sum test *p* = 2.3 × 10^-2^, FDR = 0.19) (**Supplemental Figure S8**), consistent with previous reports^5,6,22^. More evidence of significant correlations between *TERT* promoter mutations and increased expression may have been expected, but only a subset of samples with *TERT* mutations have expression data. In addition, low sequencing coverage in promoter regions limits the power of this analysis. The PCAWG drivers analysis investigated this issue specifically for two hotspot mutations in *TERT*, estimating that 216 mutations in these sites were likely not called^7^ in comparison to a total of 97 samples with *TERT* promoter mutations (71 samples with expression data).

Four other PID-N genes were found to have significant *in cis* regulatory correlations: *TP53*, *TLE4*, *TCF4*, and *DUSP22* (**Figure 3, Supplemental Figure S10**). *TP53* shows significantly reduced expression (Wilcoxon rank-sum test *p* = 1.0 × 10^-3^; FDR = 8.7 × 10^-2^) across 6 tumors with *TP53* promoter mutations from six different tumor types (**FIgure 3A, Supplemental Figure S10**). The under-expression of mutated samples is consistent with *TP53’*s well known role as a tumor suppressor gene, and links between *TP53* promoter methylation and expression have been investigated^23^. This expression change was also described by the PCAWG single-element driver discovery study^7^. *TLE4* shows significantly reduced expression in three Liver-HCC tumors (Wilcoxon rank-sum test *p* = 1.7 × 10^-2^; FDR = 0.20) with *TLE4* promoter mutations (**FIgure 3B, Supplemental Figure S10**). *TLE4* is a transcriptional co-repressor that binds to several transcription factors^24^, and *TLE4* functions as a tumor suppressor gene in acute myeloid lymphoma through its interactions with Wnt signaling^25^. Furthermore, in an acute myeloid lymphoma cell line, *TLE4* knockdown increased cell division rates while forced *TLE4* expression induced apoptosis^26^. However, the role of *TLE4* in solid tumors is not as well understood. *TCF4* shows significantly reduced expression in three Lung-SCC tumors (Wilcoxon rank-sum test *p* = 3.4 × 10^-2^; FDR = 0.27) with *TCF4* promoter mutations (**FIgure 3C, Supplemental Figure S10**). Part of the TCF4/β-catenin complex, *TCF4* encodes a transcription factor that is downstream of the Wnt signaling pathway, and low *TCF4* expression has been observed in Lung-SCC tumors^27^. *DUSP22* is significantly under-expressed in five Lung-AdenoCA patients (Wilcoxon rank-sum test *p* = 6.3 × 10^-3^; FDR = 0.024) with *DUSP22* 3’ UTR mutations and significantly over-expressed in 3 Lung-AdenoCA patients (Wilcoxon rank-sum test *p* = 7.8 × 10^-4^; FDR = 0.075) with *DUSP22* 5’ UTR mutations. These UTR mutations were mutually exclusive, and we find no support for opposing *in cis* effects in these regions. *DUSP22* encodes a phosphatase signalling protein and was recently proposed to be a tumor suppressor in lymphoma^28^.

These analyses provide additional support for a subset of PID-N genes. The small number of PID-N genes with associated gene expression changes is explained by the low number of samples with mutations in PID-N genes, the uneven availability of expression data across the tumor types, and issues of reduced coverage in non-coding regions of the genome, which may decrease the number of mutated samples and limit the ability to detect rare non-coding variants.

### The modular organization of genes impacted by coding and non-coding mutations

We identified specific protein-protein interaction subnetworks and biological pathways that were altered by coding mutations, non-coding mutations, or a combination of both types of mutations. We found significantly more interactions between PID-C genes that expected by chance using a node-degree preserving permutation test (64 interactions observed vs. 40 interactions expected, *p* < 10^-6^), a near significant number of interactions between PID-N genes (18 vs. 12 expected, *p* = 6.8 × 10^-2^), and significantly more interactions between both PID-C and PID-N genes (67 vs. 40 expected, *p* = 6 × 10^-4^), demonstrating an interplay between coding and non-coding mutations on physical protein-protein interaction networks (***Network annotation* in Methods**). Overall, we organized the interactions between PID-C and PID-N genes into five biological processes: core drivers, chromatin organization, cell proliferation, development, and RNA splicing (**Figure 4A**). While the high frequency of molecular interactions between PID-C and PID-N genes is expected since such interactions were used as a signal in pathway and network methods, the specific structure of these interactions illustrates the relative contributions of coding and non-coding mutations in individual subnetworks.

**Figure 4:**
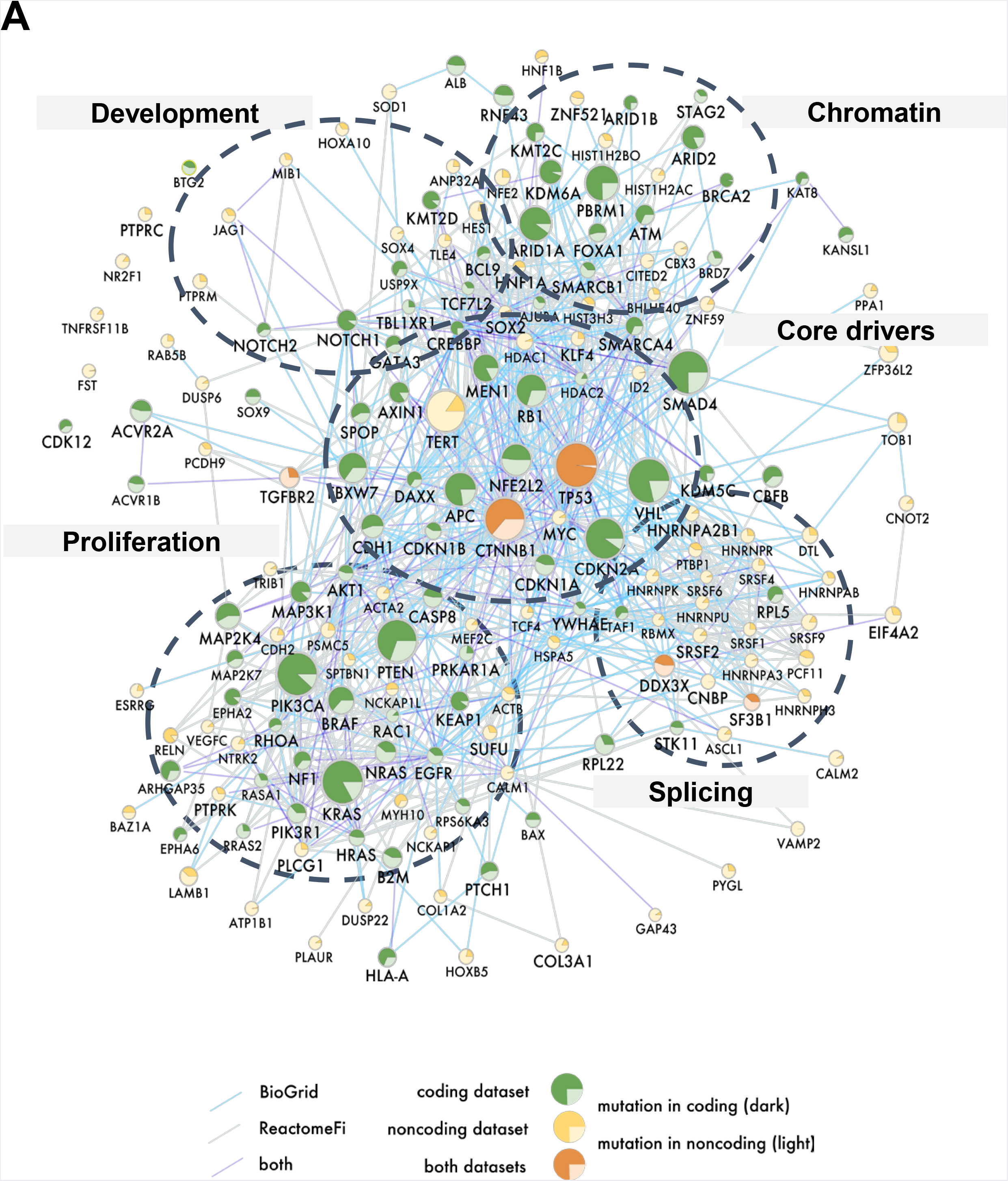
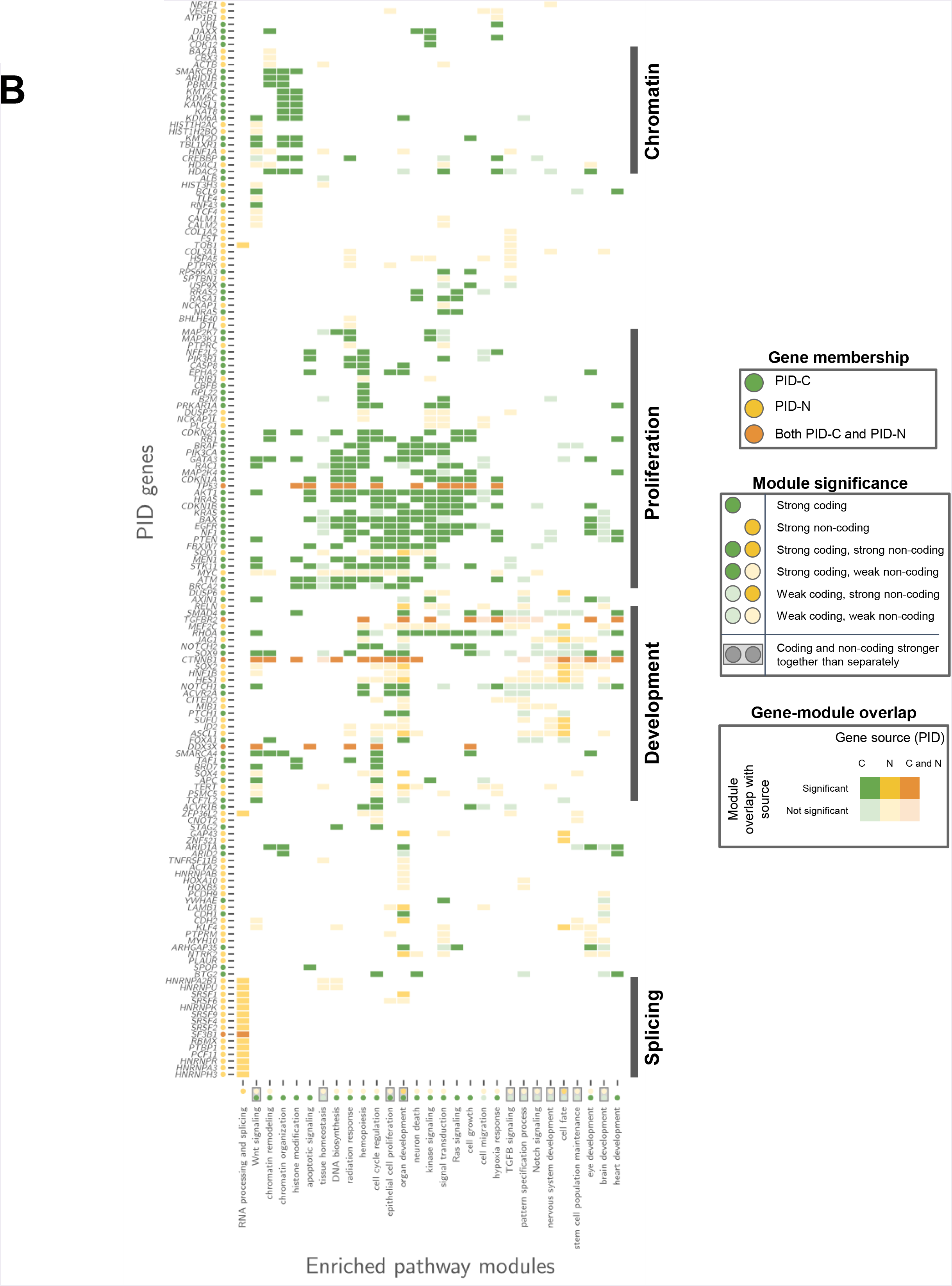
Pathway and network modules containing PID-C and PID-N genes. **(A) Network of functional interactions between PID-C and PID-N genes**. Nodes represent PID-C and PID-N genes and edges show functional interactions from the ReactomeFI network (grey), physical protein-protein interactions from the BioGRID network (blue), or interactions recorded in both networks (purple). Node color indicates PID-C genes (green), PID-N genes (yellow), or both PID-C and PID-N genes (orange);node size is proportional to the score of the corresponding gene; and the pie chart diagram in each node represents the relative proportions of coding and non-coding cancer mutations associated with the corresponding gene. Dotted outlines indicate clusters of genes with roles in chromatin organization and cell proliferation, which predominantly contain PID-C genes; development, which includes comparable amounts of PID-C and PID-N genes; and RNA splicing, which contains PID-N genes. A core cluster of genes with many known drivers are also indicated. **(B) Pathway modules containing PID-C and PID-N genes**. Each row in the matrix corresponds to a PID-C or PID-N gene, and each column in the matrix corresponds to a pathway module enriched in PID-C and/or PID-N genes (see **Methods**). A filled entry indicates a gene (row) that belongs to one or more pathways (column) colored according to gene membership in PID-C genes (green), PID-N genes (yellow), or both PID-C and PID-N genes (orange). A darkly colored entry indicates that a PID-C or PID-N gene belongs to a pathway that is significantly enriched for PID-C or PID-N genes, respectively. A lightly colored entry indicates that a PID-C or PID-N gene belongs to a pathway that is significantly enriched for the union of PID-C and PID-N genes but not for PID-C or PID-N genes separately. Enrichments are summarized by circles adjacent each pathway module name and PID gene name. Boxed circles indicate that a pathway module contains a pathway that is significantly more enriched for the union of the PID-C and PID-N genes than the PID-C and PID-N results separately. The enriched modules and PID genes are clustered into four biological processes: chromatin, development, proliferation, and RNA splicing as indicated, with differing contributions of PID-C and PID-N genes.

We further characterized the molecular pathways enriched among our PID-C and PID-N using the g:Profiler web server^29^ (**Figure 4B, Supplemental Figure S9, Supplemental Tables S15-S18, *Pathway annotation* in Methods**). Since our methods use pathway databases and interaction networks as prior knowledge, enrichment with known pathways is expected. However, the enrichment results provide clues about the modular organization of the pathways and allow us to assess the relative contributions of coding and non-coding mutations in each pathway. Overall, 63 pathways were enriched for PID-C genes and 13 pathways were enriched for PID-N genes (FDR < 10^-6^).

We further grouped these pathways into 29 modules using overlaps between annotated pathways recorded in the pathway enrichment map (**Supplemental Figure S11**). For each enriched module, we examined whether PID-C, PID-N, or both types of genes were responsible for the observed enrichment. This produced a clustering of modules and PID genes into four biological processes: chromatin organization, cell proliferation, development and RNA splicing (**Figure 4B**).

We found that pathways in the chromatin and cell proliferation processes — including chromatin remodeling and organization, histone modification, apoptotic signaling, signal transduction, Ras signaling, and cell growth — were altered primarily by coding mutations in PID-C genes. This is not surprising as these pathways contain many well-known cancer genes, such as *TP53, KRAS, BRAF*, cyclin dependent kinase inhibitors, *EGFR, PTEN*, and *RB1*.

Several signaling pathways contain significant numbers of both PID-C and PID-N genes, indicating that non-coding mutations provide additional avenues for disrupting key molecular interactions. These pathways include the Wnt signaling pathway (FDR = 6.8 × 10^-13^), which was predominantly targeted by coding mutations but was also targeted by non-coding mutations in several PID-N genes, including *TERT* (103 mutations), *HNF1A/B* (24 mutations), *TLE4* (32 mutations), *TCF4* (93 mutations), and *CTNNB1* (17 mutations) (**Supplemental Figure S12A**). The Notch signaling pathway (FDR = 6.8 × 10^-7^) was associated with comparable numbers of PID-C and PID-N genes, including the PID-N genes *JAG1* and *MIB1* that encode ligands and the PID-N transcription factors *ACL1*, *HES1*, and *HNF1B* (66 non-coding mutations in total) (**Supplemental Figure S12B**). The TGF-β signaling pathway (FDR = 3.2 × 10^-7^) also contained both PID-C and PID-N genes, including the PID-N genes *HES1*, *HNF1A/B*, *HSPA5, MEF2C* as well as *TGFBR2* and *CTNNB1* (214 coding mutations and 166 non-coding mutations), which are both PID-C and PID-N genes.

We found that several developmental processes were altered by significant numbers of both PID-C and PID-N genes. Cell fate determination (FDR = 2.0 × 10^-7^) was predominantly affected by non-coding mutations in the PID-N genes *DUSP6, MEF2C, JAG1*, *SOX2, HES1*, *ACL1*, *ID2, SUFU, and KLF4* (total 191 non-coding mutations) but also includes PID-C genes *BRAF, GATA3, NOTCH1/2*. Pathways related to nervous system development (FDR = 5.8 × 10^-8^) were enriched for the PID-N genes *ASCL1*, *CTNNB1*, *ID2, SUFU*, and *TERT* that have known roles in cancer^30,31^, complementing the PID-C genes *NOTCH1*, *PTEN* and *RHOA* that also have known cancer roles. The pattern specification process (FDR = 8.8 × 10^-8^) was also affected by both coding and non-coding mutations, including the PID-N genes *ASCL1*, *SUFU*, and *RELN* and the PID-C genes *ATM* and *SMAD4*. In these cases, non-coding mutations complement coding mutations that disrupt these pathways, covering significant numbers of additional patients.

Intriguingly, we find that RNA splicing pathways were affected primarily by non-coding mutations (FDR = 7.6 × 10^-9^). A total of 17 PID-N genes belonged to splicing-related pathways (**Supplemental Figure S12C**), including several heterogeneous nuclear ribonucleoproteins (hnNRP) and serine and arginine rich splicing factors (SRSFs). None of these PID-N genes were significantly mutated according to single-element tests of the PCAWG driver discovery analysis. We did not find any significant (FDR < 0.1) *in cis* associations between non-coding mutations and altered expression of these genes. Thus, we explored potential *in trans* effects on pathway expression changes. We found that non-coding mutations in splicing-related PID-N genes largely recapitulate a recently published association by TCGA^32^ between coding mutations in several splicing factors and differential expression of 47 pathways (**Figure 5**). Specifically, we identified three clusters of mutations (C1, C2, and C3 in **Figure 5A and Figure 5B**) from our differential expression analysis. Each of these clusters contained at least one coding mutation in the splicing genes *SF3B1*, *FUBP1*, and *RBM10* as reported in ^32^, with non-coding mutations in splicing-related PID-N genes showing similar gene expression signatures. The joint analysis of coding and non-coding mutations in splicing factors also recovered the two groups of enriched pathways (P1 and P2 in **Figure 5A, Supplemental Figure S13**) reported in ^32^. One group (P1) is characterized by immune cell signatures and the other group (P2) reflects mostly cell-autonomous gene signatures of cell cycle, DDR, and essential cellular machineries^32^. The similarity between the gene expression signatures for non-coding mutations in several PID-N splicing factors and coding mutations in splicing factor genes^32^ supports a functional role for splicing-related PID-N genes in altering similar gene expression programs.

**Figure 5:**
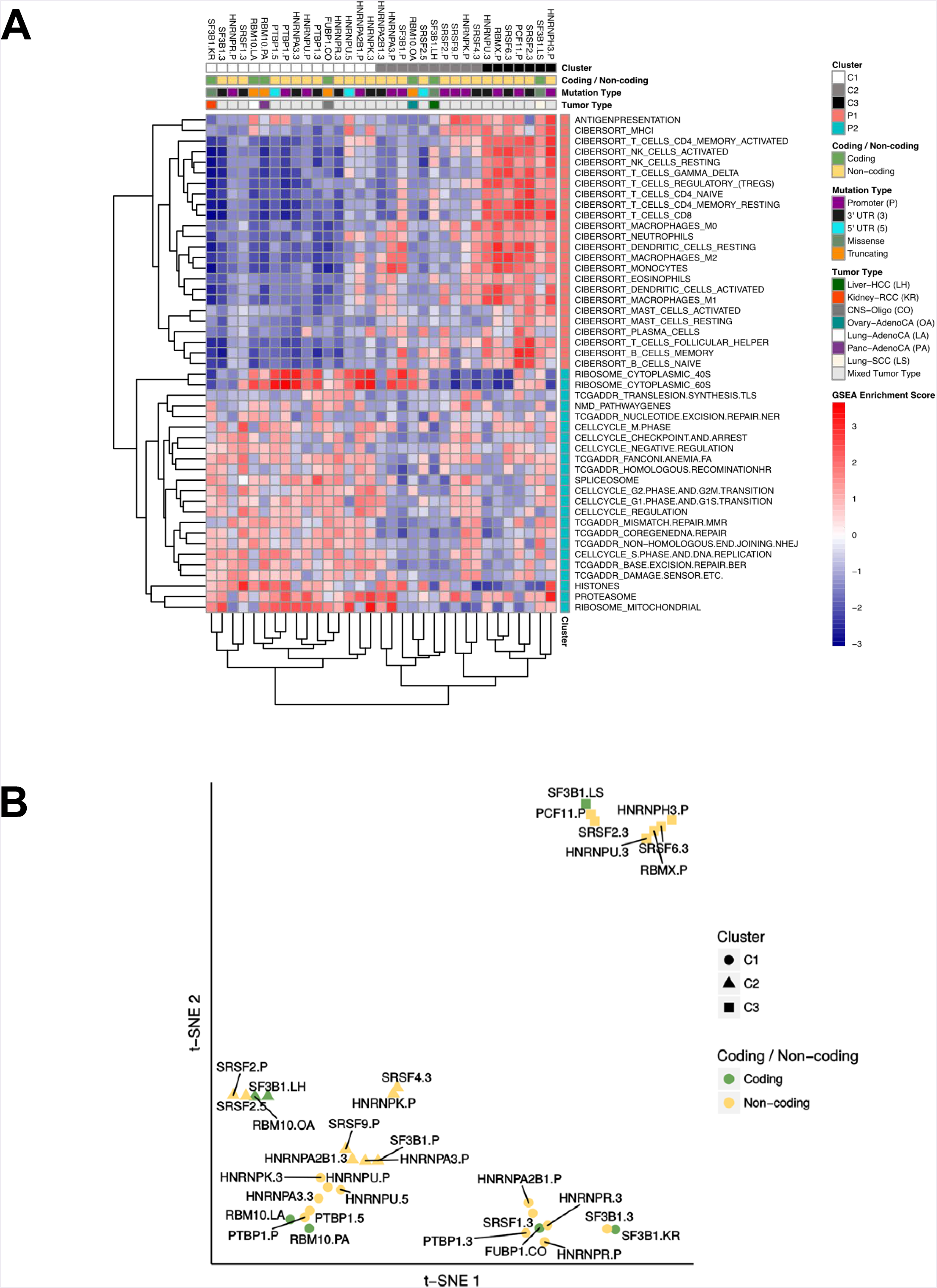
RNA splicing factors are targeted primarily by non-coding mutations and alter expression of similar pathways as coding mutations in splicing factors. **(A) Heatmap of Gene Set Enrichment Analysis (GSEA) Normalized Enrichment Scores (NES)**. The columns of the matrix indicate non-coding mutations in splicing-related PID-N genes and coding mutations in splicing genes reported in ^32^ and the rows of the matrix indicate 47 curated gene sets^32^. Red heatmap entries represent an upregulation of the pathway in the mutant samples with respect to the non-mutant samples and blue heatmap entries represent a downregulation. The first column annotation indicates mutation cluster membership according to common pathway regulation. The second column annotation indicates whether a mutation is a non-coding mutation in a PID-N gene or a coding mutation^32^, with the third column annotation specifies the aberration type (promoter, 5’ UTR, 3’ UTR, missense, or truncating). The fourth column annotation indicates the cancer type for coding mutations from ^32^. The mutations cluster into 3 groups: C1, C2, and C3. The pathways cluster into two groups^32^: P1 and P2, where P1 contains an immune signature gene sets and P2 contains cell autonomous gene sets as reported in ^32^. **(B) tSNE plot of mutated elements illustrates clustering of gene expression signatures for samples with non-coding mutations in splicing-related PID-N genes with gene expression signatures for coding mutations in previously published splicing factors**. The shape of each point denotes the mutation cluster assignment (C1, C2, or C3), and the color represents whether the corresponding gene is a PID-N gene with non-coding mutations or a splicing factor gene with coding mutations^32^.

In addition to the above modules, we also found that transcription factors were well represented among both the PID-C and PID-N genes. In total, 9 PID-C genes are transcription factors (*ARHGAP35, ARID2, FOXA1*, *GATA3*, *NFE2L2, SMAD4*, *SOX9*, *TCF7L2, TP53;* FDR = 2.1 × 10^-10^), while 19 PID-N genes are transcription factors (*ASCL1*, *BHLHE40, ESRRG*, *HES1*, *HNF1A*, *HNF1B*, *HOXA10, HOXB5, KLF4*, *MEF2C*, *MYC*, *NFE2*, *NR2F1*, *SOX2, SOX4*, *TCF4*, *TP53, ZNF521*, *ZNF595;* FDR = 4.1 × 10^-20^).

## Discussion

While single-region tests in the PCAWG project identified only a few non-coding driver elements, our integrative pathway and network analysis further expands the list of genes with possible non-coding driver mutations, extending into the “long tail” of rare mutations. In particular, we find that genes with either coding or non-coding mutations are linked in pathways and networks, and that pathway databases and interaction networks can be leveraged as prior knowledge to identify additional possible non-coding drivers that are too infrequently mutated to be detected by single-element tests. In total, our integrative pathway analysis identified 87 pathway-implicated driver genes with coding variants (PID-C) and 93 pathway-implicated driver genes with non-coding variants (PID-N). Importantly, 90 PID-N genes were not statistically significant (FDR > 0.1) by single-element tests on non-coding mutation data, and these genes are key candidates for future experimental characterization. Among them, we find that promoter mutations in *TP53, TLE4*, and *TCF4* are associated with reduced expression of these genes.

We find that coding and non-coding driver mutations largely target different genes, and contribute differentially to pathways and networks perturbed in cancer. While some cancer pathways are targeted by both coding and non-coding mutations, such as the Wnt and Notch signaling pathways, other pathways appear to be predominately altered by one class of mutations. In particular, we find non-coding mutations in multiple genes in the RNA splicing pathway, and samples with these mutations exhibit gene expression signatures that are concordant with gene expression changes observed in samples with coding mutations splicing factors *SF3B1*, *FUBP1*, and *RBM10^32^*. Together these results demonstrate that rare non-coding mutations may result in similar perturbations to both common and complementary biological processes.

There are several caveats to the results reported in this study. First, there is relatively low power to detect non-coding mutations in the cohort, particularly in cancer types with small numbers of patients. Second, transcriptomic data was available for only a subset of samples, further reducing our ability to validate our predictions using gene expression data. Third, our pathway and network analysis relied on the driver *p*-values from the PCAWG consensus driver analysis^7^. This analysis accounts for regional variations in the background mutation rate across the genome. However, if these corrections are inadequate and the uncorrected confounding variables are correlated with gene membership in pathways and subnetworks, then the false positive rates in our analysis may be higher than estimated. All of these factors, plus other unknown confounding variables, make it difficult to assess the false discovery rate of our predictions, particularly for PID-N genes. Further experimental validation of these predictions is necessary to determine the true positives from false positives in our PID gene lists.

While pathway and network analysis was successful in revealing potential new cancer-associated genes impacted by non-coding mutations, future investigations that consider the changing landscape of gene regulation and pathway interactions across tissues may offer a new perspective on the data. Specifically, each cell type has a different epigenetic wiring and regulatory machinery, and non-coding mutations may target cell type-specific vulnerabilities. Approaches that incorporate tissue-specific gene-gene regulatory logic may be successful in revealing new classes of drivers unexplored with our current approaches.

In conclusion, our pathway- and network-driven strategies enable us to interpret the coding and non-coding landscape ofxs tumor genomes to discover driver mechanisms in interconnected systems of genes. This approach has multiple benefits. First, by broadening our mutation analysis from single genomic elements to pathways and networks of multiple genes, we identify new components of known cancer pathways that are recurrently altered by both coding and non-coding mutations, and thus likely to be important in cancer. Second, we identify new pathways and subnetworks that would remain unseen in an analysis focusing on coding sequences. Investigation of the coding and non-coding mutations that perturb these pathways and networks will enable more accurate patient-stratification strategies, pathway-focused biomarkers, and therapeutic approaches.

## Acknowledgements

B.J.R. received funding from NIH grants U24CA211000 and R01HG007069. J.M.S. received funding from NIH grants U24CA143858, R01CA180778, and U24CA210990. J.R. received funding from the Ontario Institute for Cancer Research (OICR) Investigator Award provided by the Government of Ontario, Operating Grant from Cancer Research Society (CRS) (#21089), and the Natural Sciences and Engineering Research Council of Canada (NSERC) Discovery Grant (#RGPIN-2016-06485). K.M. received funding from IWT/SBO NEMOA and FWO 3G046318 and G.0371.06 grants. J.M.G.I. received funding from from the Novo Nordisk Foundation (NNF17OC0027594 and NNF14CC0001) and the Innovation Fund Denmark (5184-00102B). S.B. received funding from the Novo Nordisk Foundation (NNF17OC0027594 and NNF14CC0001). J.B. received funding from the BioTalent Canada Student Internship Program. A.V. and M.V. received funding from the Joint BSC-IRB-CRG Program in Computational Biology and the Severo Ochoa Award (SEV 2015-0493).

We thank Esther Rheinbay and the rest of the PCAWG 2-5-9-14 group for their assistance with PCAWG consensus driver analysis data and Angela Brooks for her help with splicing analysis.

## Supplemental Figure Legends

**Figure S1:**
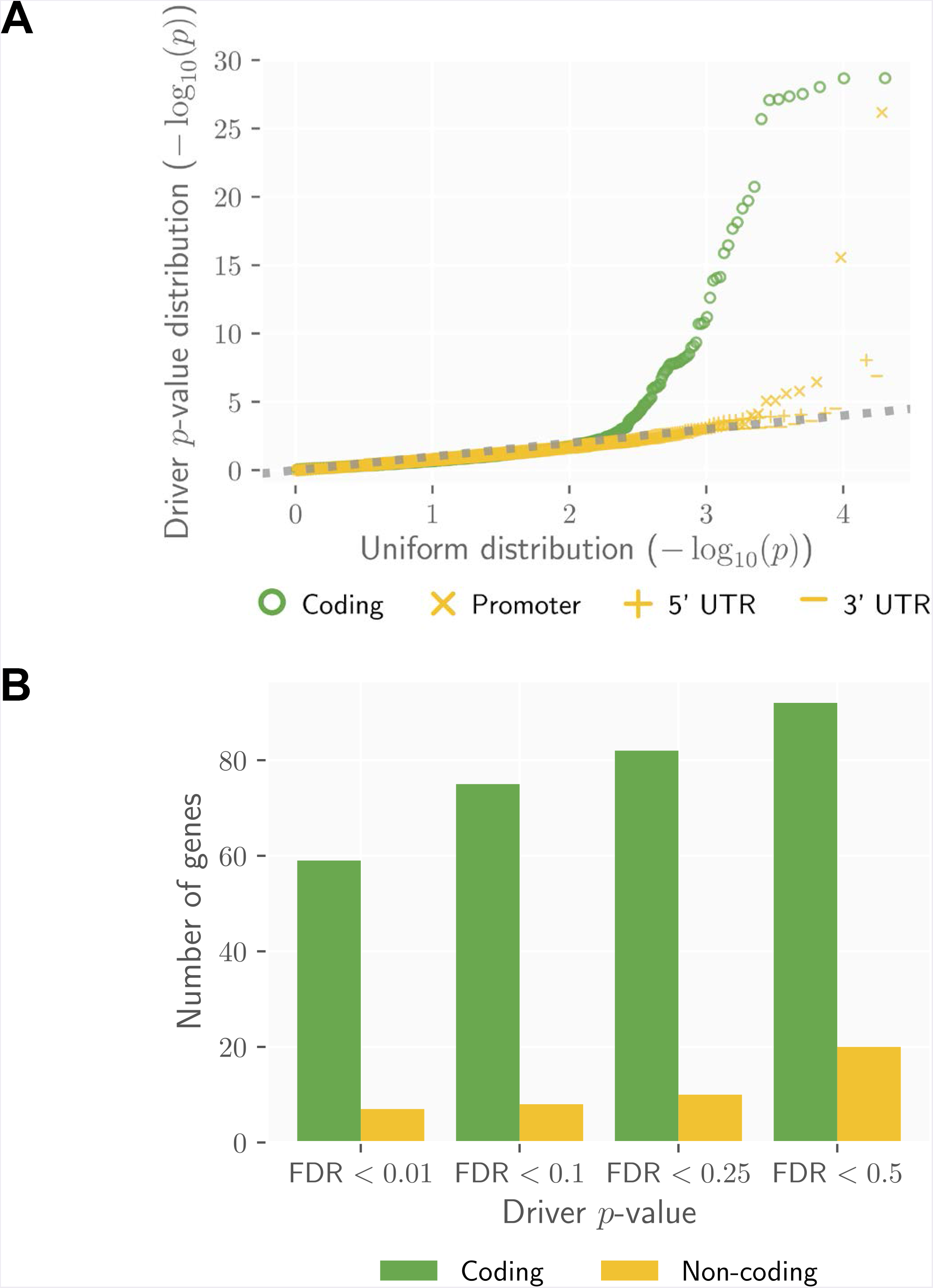
Driver *p*-value distributions for coding and non-coding regions of the genome. **(A)** Distribution of driver *p*-values from single-element tests on coding and non-coding (promoter, 5’ UTR, 3’ UTR) regions of the genome. **(B)** Numbers of genes with driver *p*-values from single-element tests with q-values satisfying *q* < 0.01, 0.1, 0.25, 0.5 on coding and non-coding (promoter, 5’ UTR, 3’ UTR) regions of the genome.

**Figure S2:**
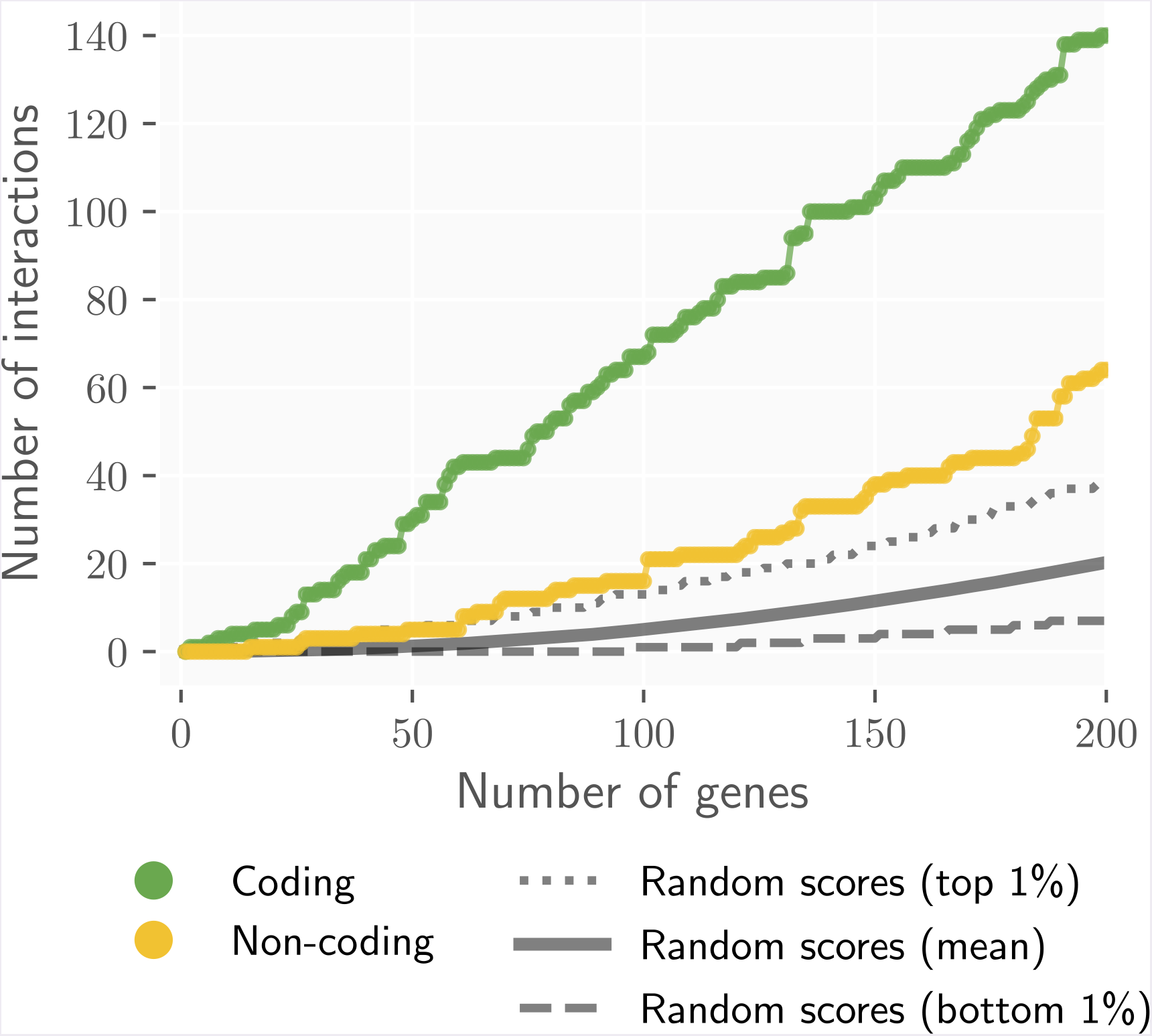
Statistically significant network interactions between genes with highest driver *p*-values. Genes in the BioGRID high-confidence functional interaction network with the highest coding and non-coding (promoter, 5’ UTR, 3’ UTR) driver *p*-values have statistically significant numbers of interactions compared to genes chosen uniformly at random from the network. We rank network genes by their coding or non-coding driver *p*-values (by single element q-values) and show the number of interactions between the genes with highest observed coding (green) and non-coding (yellow) *p*-values as well as random (gray) *p*-values using 1,000 permutations among network genes.

**Figure S3:**
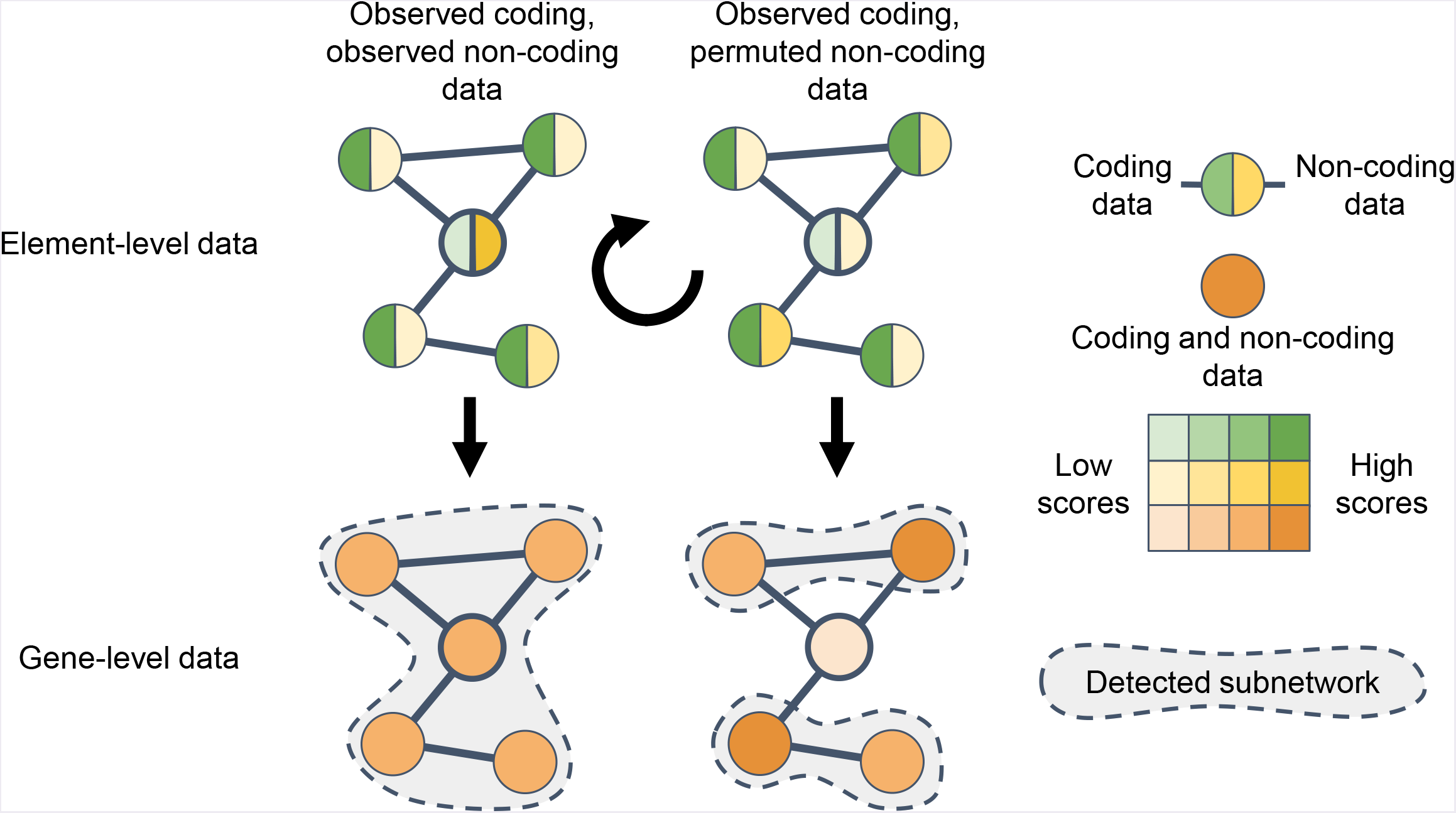
Illustration of non-coding value-added (NCVA) procedure. **Illustration of the NCVA procedure for identifying results on coding and non-coding data with with significant contributions from non-coding data.** (A) Top left: The central gene has a high non-coding gene score, and the four neighboring genes have high coding gene scores. **(B)** Bottom left: All five genes have strong combined coding and non-coding gene scores. A pathway/network method identifies a subnetwork of all five genes. **(C)** Top right: After preserving coding gene scores and permuting non-coding gene scores, the central gene has a low non-coding gene score, and the four neighboring genes still have high coding gene scores. **(D)** Bottom right: Four of the five genes have strong combined coding and non-coding gene scores, but the central gene does not. A pathway/network method identifies two subnetworks of two genes, excluding the central gene, which becomes a potential NCVA gene If a gene identified by a pathway/network method using observed coding and observed non-coding gene scores and consistently omitted (*p* < 0.1) by the method using observed coding scores and permuted non-coding gene scores, then we identify it as a non-coding value-added (NCVA) gene for that method because the non-coding data makes a significant contribution to that gene’s discovery by a method on coding and non-coding data.

**Figure S4:**
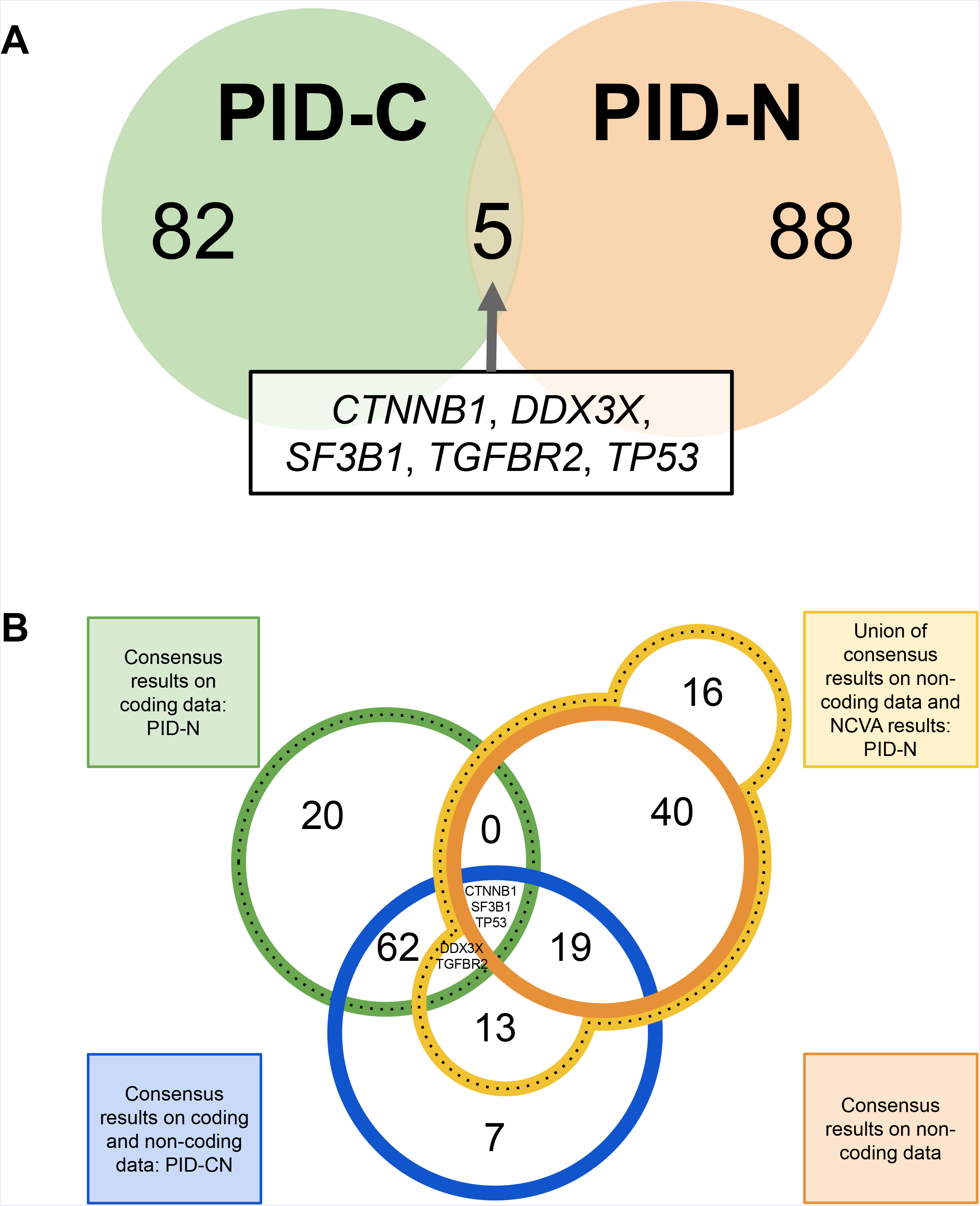
Overlap of consensus results for pathway and network methods. **(A) PID-C and PID-N genes have negligible overlap**. Only 5 genes (*CTNNB1, DDX3X, SF3B1, TGFBR2, TP53* are both PID-C and PID-N genes. **(B) Overlap of all consensus results**. Four-circle Venn diagram for the overlap of the consensus results on coding data, i.e., PID-C genes; consensus pathway/network results on non-coding data; consensus pathway/network results on coding and non-coding data; and the union of the consensus results on non-coding data and the non-coding value-added (NCVA) results, i.e., PID-N genes.

**Figure S5.**
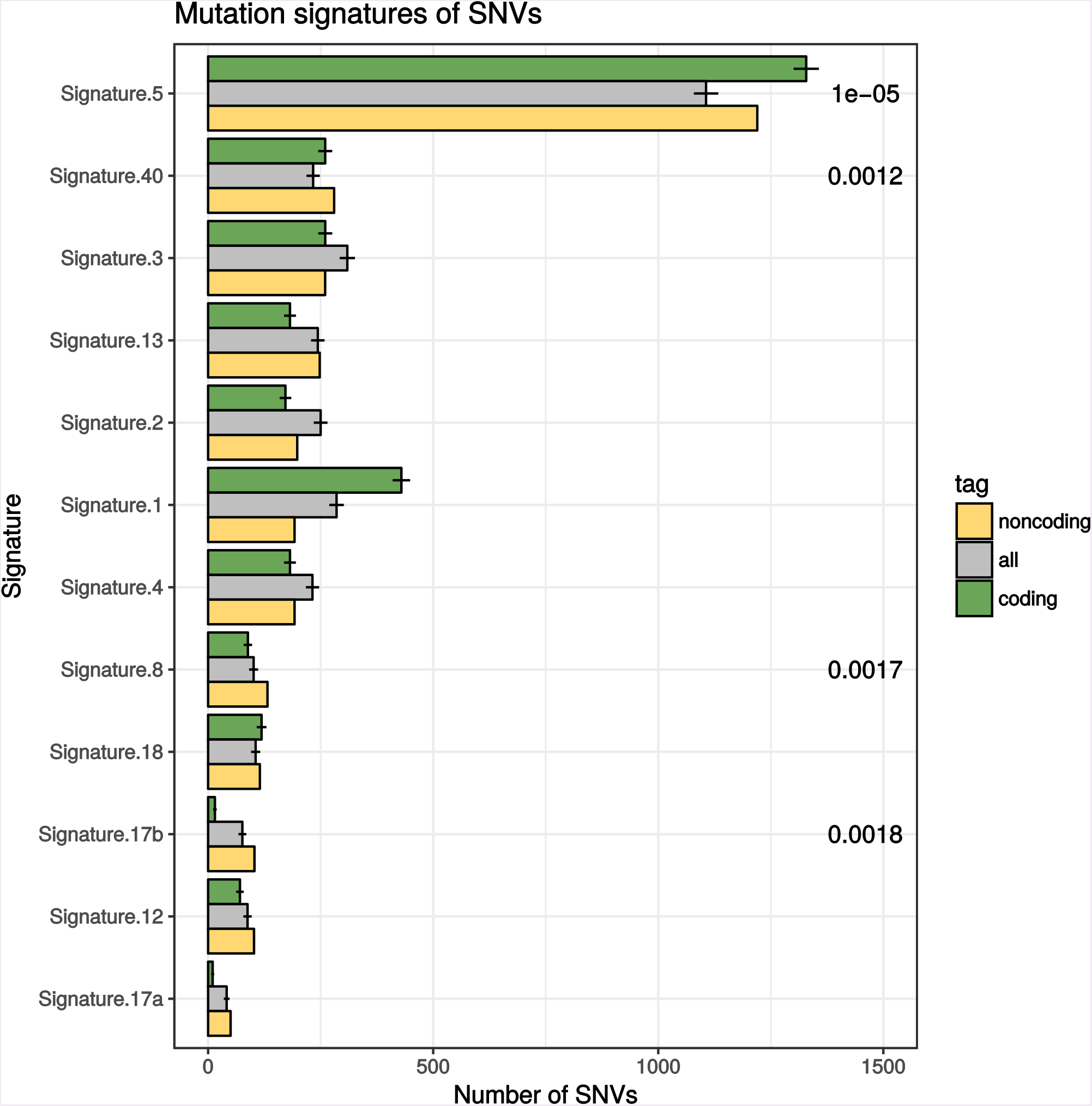
Mutation signatures of SNVs in PID-C and PID-N genes. Bar plot shows predicted mutation signatures of observed mutations among PID-N genes (yellow) compared with randomly sampled mutations in all coding and non-coding elements (grey). Mutations in PID-C genes are shown as a positive control (green). *p*-values were computed with custom permutation tests and show enrichment of mutation signatures within PID-N genes (yellow) relative to all sampled mutations (grey). *p*-values with *p* < 0.05 are shown.

**Figure S6:**
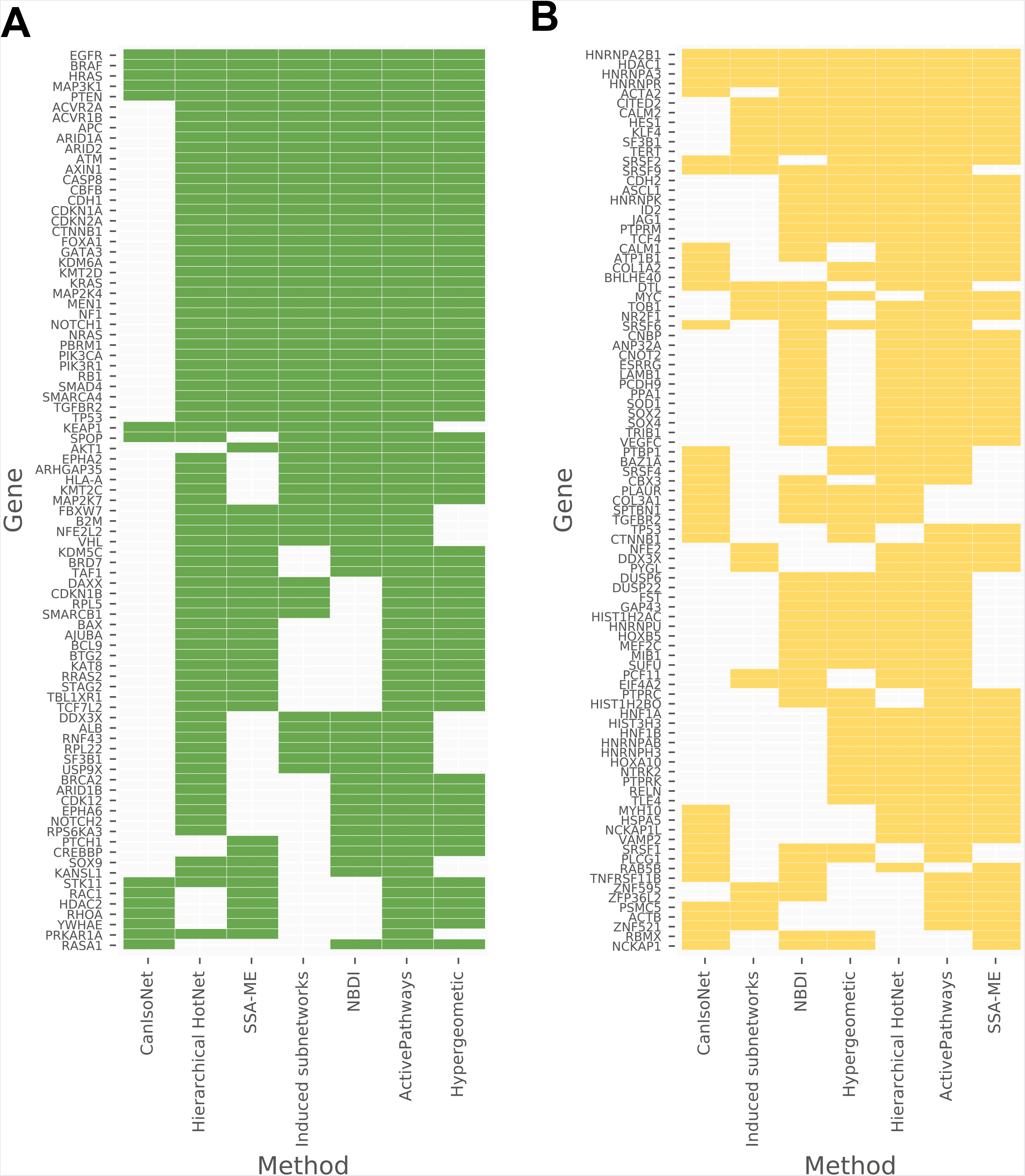
Annotations of PID-C and PID-N genes. **(A) Pathway and network method contributions to PID-C genes**. The left matix (green entries) depicts method contributions to the PID-C genes. Each row is a PID-C gene, each column is a pathway or network method, and each filled entry indicates that a method contains a PID-C gene. Both genes and methods are ordered by hierarchical clustering (Jaccard index, single-linkage clustering; hierarchies omitted) to show genes that are reported by similar methods and methods that report similar gene sets. **(B) Pathway and network method contributions to PID-N genes**. The right matrix (yellow entries) shows method contributions to the PID-N results. The matrix is similar to (A) except each filled entry indicates that a method contains a PID-N gene.

**Figure S7:**
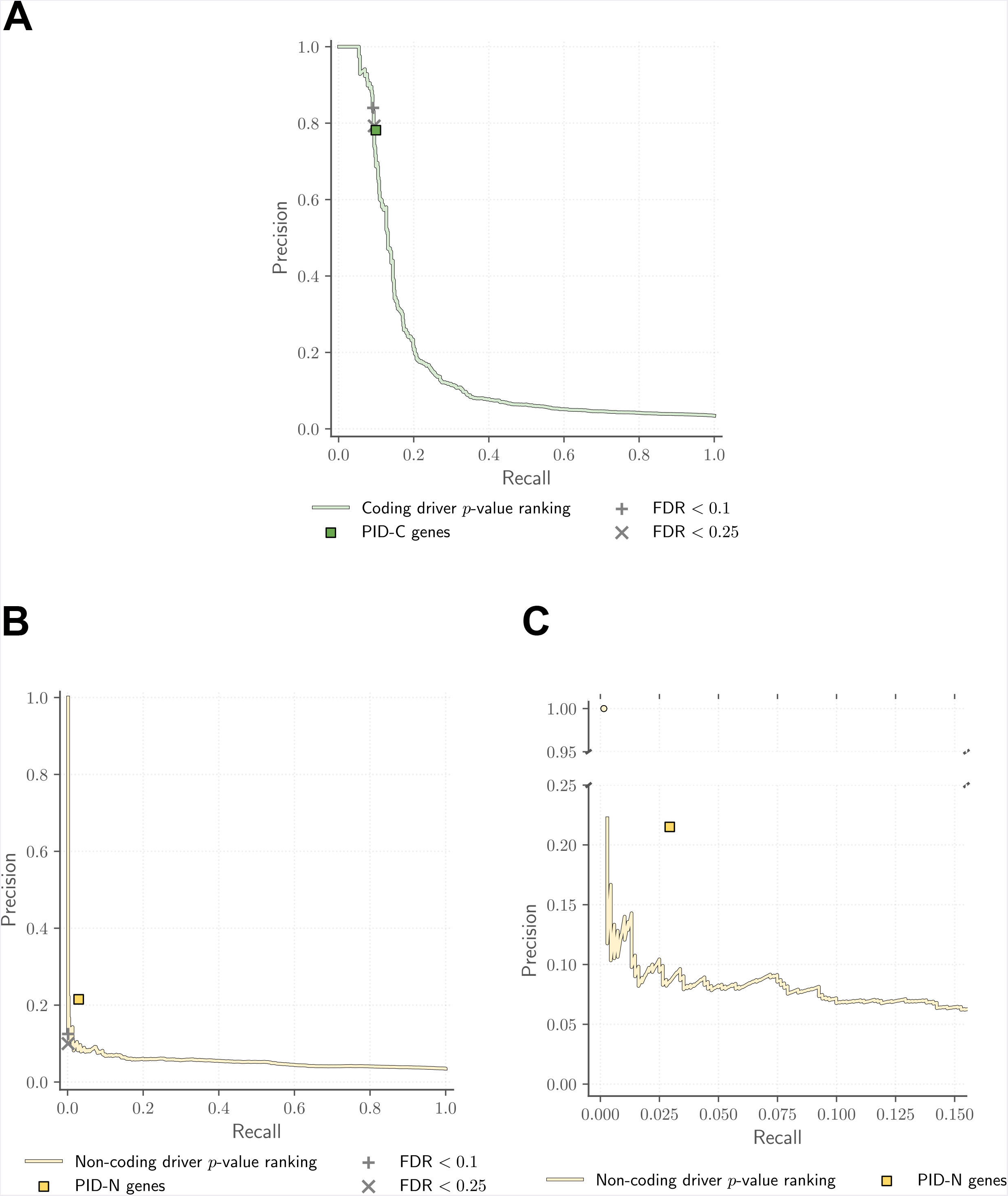
Enrichment of genes with high driver *p*-values, pathway and network method results for COSMIC Cancer Gene Census (CGC) genes. **(A) Precision and recall of coding driver *p*-values and PID-C genes for COSMIC CGC genes**. Precision and recall of genes ranked by driver *p*-values on coding elements and PID-C genes with COSMIC CGC genes. Driver FDR thresholds of 0.1 and 0.25 are highlighted. **(B, C) Precision and recall of of non-coding driver *p*-values and PID-N genes for COSMIC CGC genes**. Precision and recall of genes ranked by driver *p*-values on non-coding (promoter, 5’ UTR, 3’ UTR) and PID-N genes with COSMIC CGC genes. Driver FDR thresholds of 0.1 and 0.25 are highlighted. The left-most plot (B) shows the full y-axis, and right-most plot (C) shows a broken y-axis.

**Figure S8:**
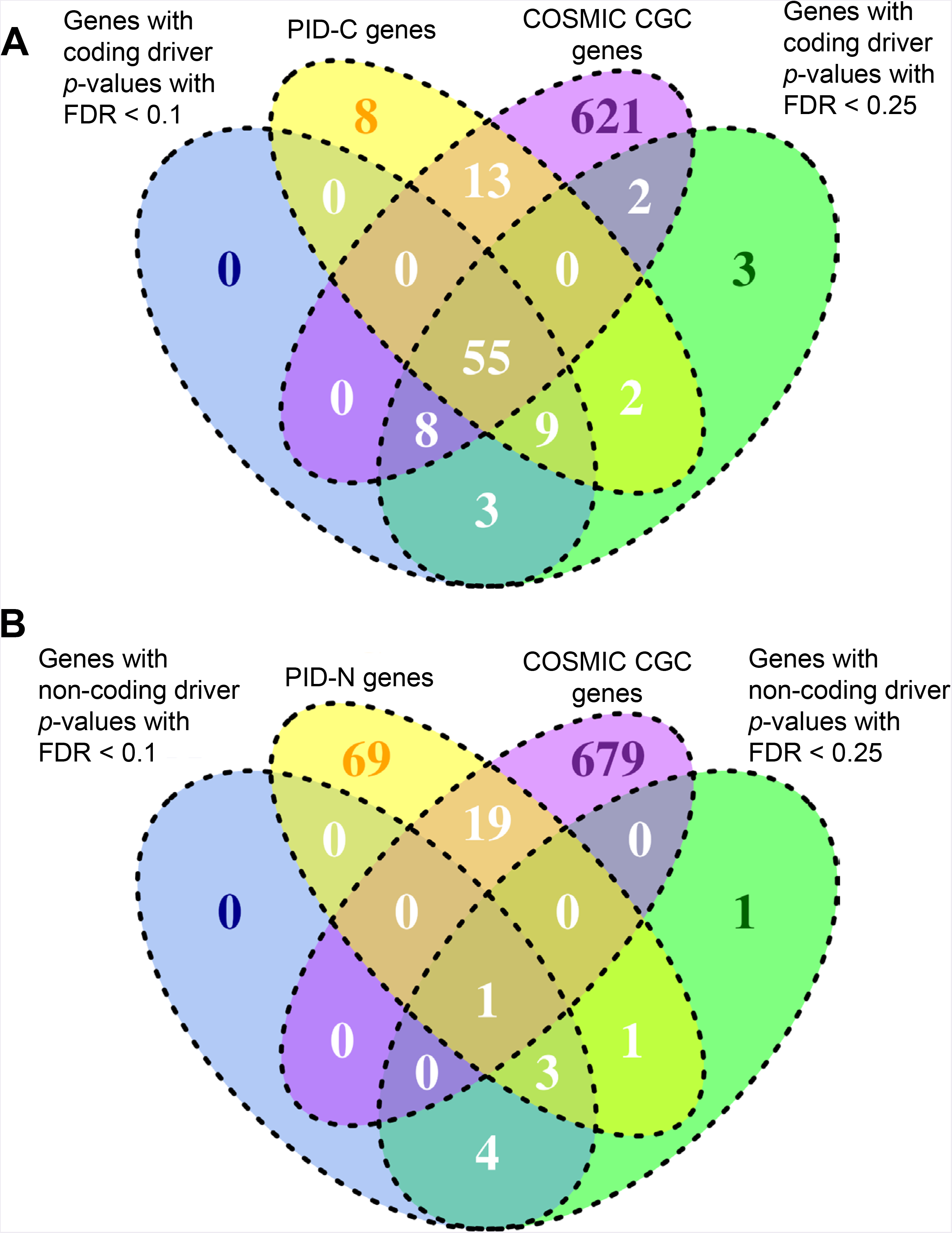
Overlap of genes with high driver *p*-values, pathway and network method results with COSMIC Cancer Gene Census (CGC) genes. **(A) Venn diagram for PID-C genes**. Overlap of genes with coding driver *p*-values with FDR < 0.1, genes with coding driver *p*-values with FDR < 0.25, PID-C genes, and COSMIC CGC genes. **(B) Venn diagram for PID-N genes**. Overlap of genes with non-coding (promoter, 5’ UTR, 3’ UTR) driver *p*-values with FDR < 0.1, genes with non-coding *p*-values with FDR < 0.25, PID-N genes, and COSMIC CGC genes.

**Figure S9:**
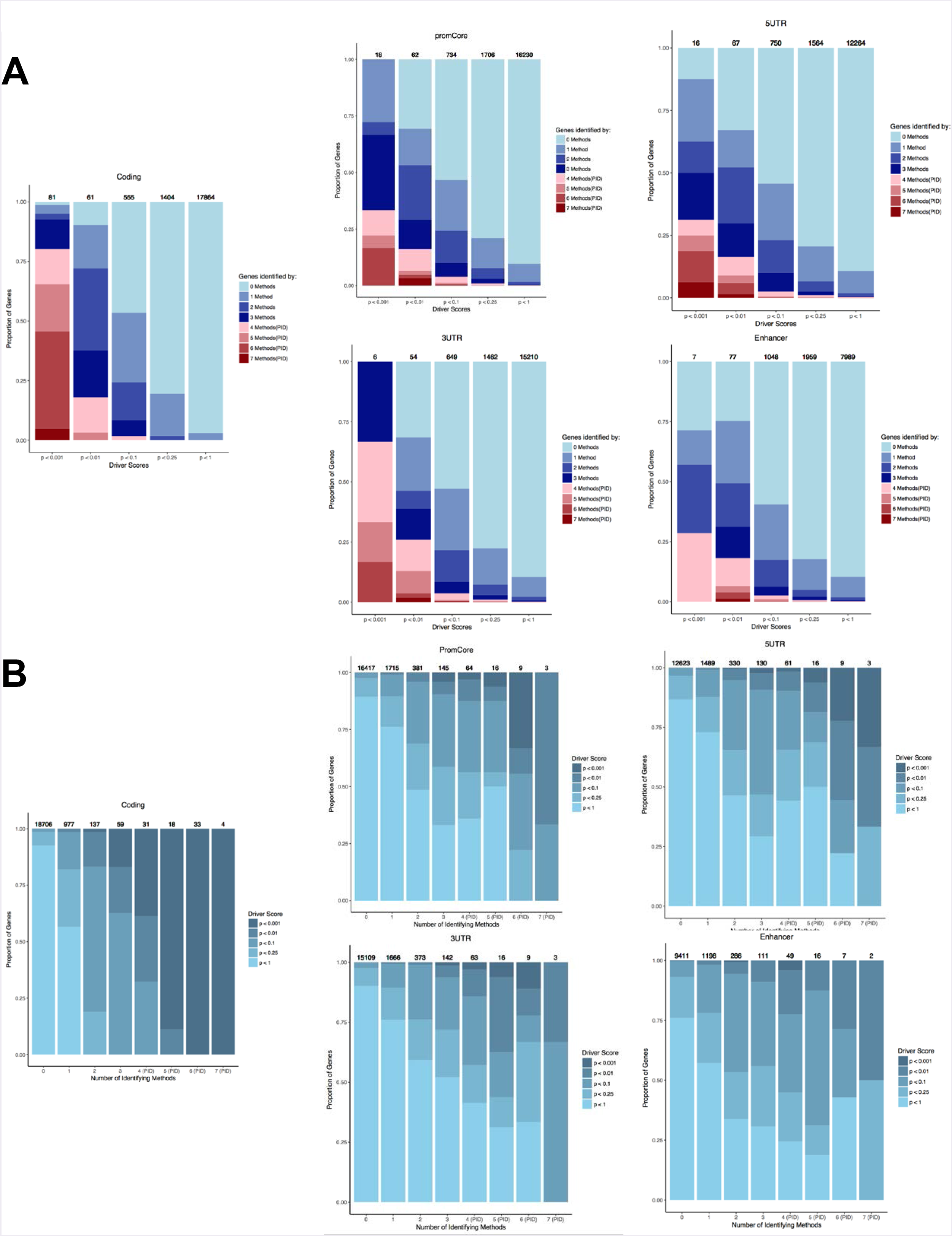
Distribution of gene scores and pathway and network method results. **(A) Driver *p*-values of genes identified by pathway and network methods**. Stacked bar chart showing distribution of coding and non-coding (promoter, 5’ UTR, 3’ UTR, enhancer) driver *p*-values for genes identified by different numbers pathway and network methods, where genes identified by a majority (≥ 4/7) of methods are PID genes. **(B) Driver *p*-values of genes identified by pathway and network methods**. Bar chart showing distribution of number of genes identified by pathway and network methods for genes with driver *p*-values with *p* < 0.001, 0.01, 0.1, 0.25, 1.

**Figure S10:**
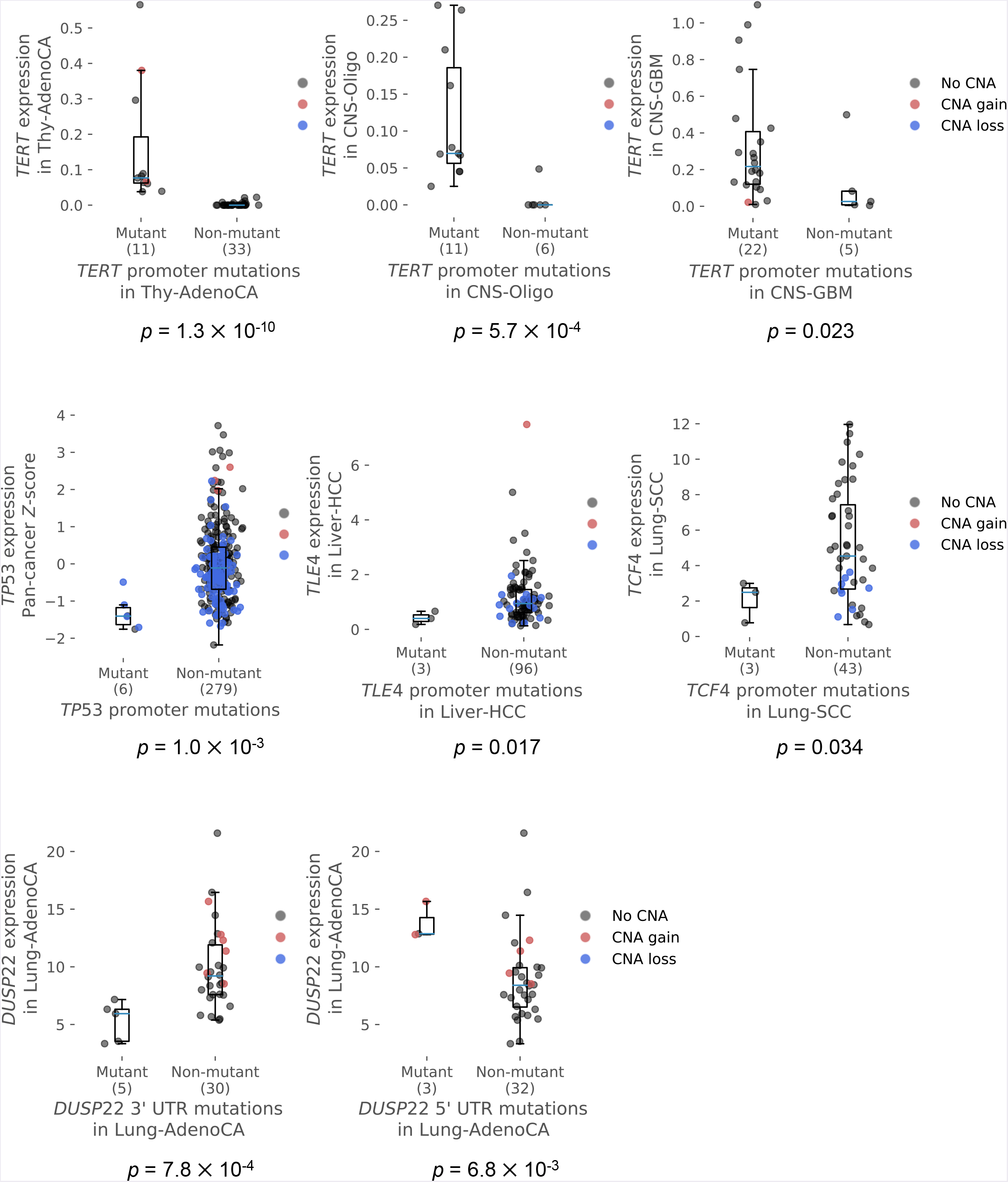
Gene expression changes are correlated with mutations in PID-N genes. All non-coding mutations in PID-N genes that show significant expression changes (rank sum FDR < 0.3): promoter mutations in *TERT, TP53, TLE4*, and *TCF4* and 3’ UTR and 5’ UTR mutations in *DUSP22*. Each plot shows the expression (FPKM-UQ values on individual tissue types or z-scores for FPKM-UQ values across tissue types) of a gene for patients with (left) and without (right) mutations in that gene, where each point in the plot indicates the expression of each patient. Copy number gains (numeric copy number gain of at least 1) and losses (numeric copy number loss of at least 1) are highlighted in red and blue, respectively.

**Figure S11:**
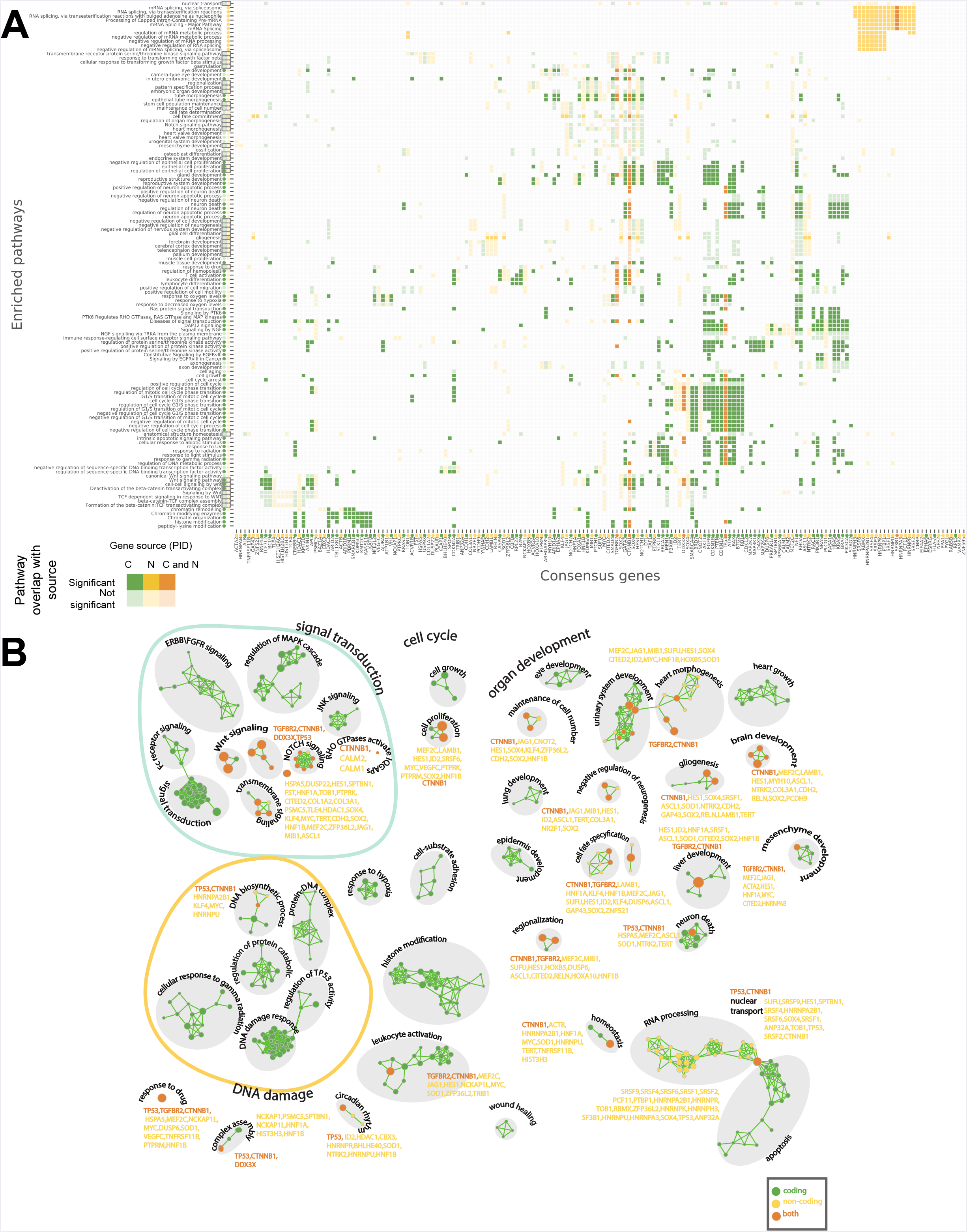
Pathways containing PID-C and PID-N genes. **(A) Pathways containing PID-C and PID-N genes**. This figure shows the pathways in the pathway modules in Figure 4B. Each row corresponds to a pathway that is enriched (see **Methods**) in PID-C and/or PID-N genes, and each column is a PID-C or PID-N gene. A filled entry in the table indicates a gene (column) that belongs to a pathway (row), colored according to PID-C genes (green), PID-N genes (yellow), or both (orange). Dark colors indicate that the corresponding module contains a pathway that is significantly enriched (dark) or not (light) for) that include the corresponding gene. Enrichments are summarized by circles adjacent each pathway name and consensus gene name. Boxed circles indicate that a pathway contains a pathway that is significantly more enriched for the union of the PID-C and PID-N than the PID-C and PID-N results separately. **(B) Pathway enrichment map**. Nodes in the enrichment map represent pathways, and edges indicate highly overlapping pathways. Node color shows if detected pathways are supported (high pathway enrichment) by the PID-C gene set (green), PID-N gene set (yellow) or both (orange). Node size indicates number of genes in the pathway.

**Figure S12:**
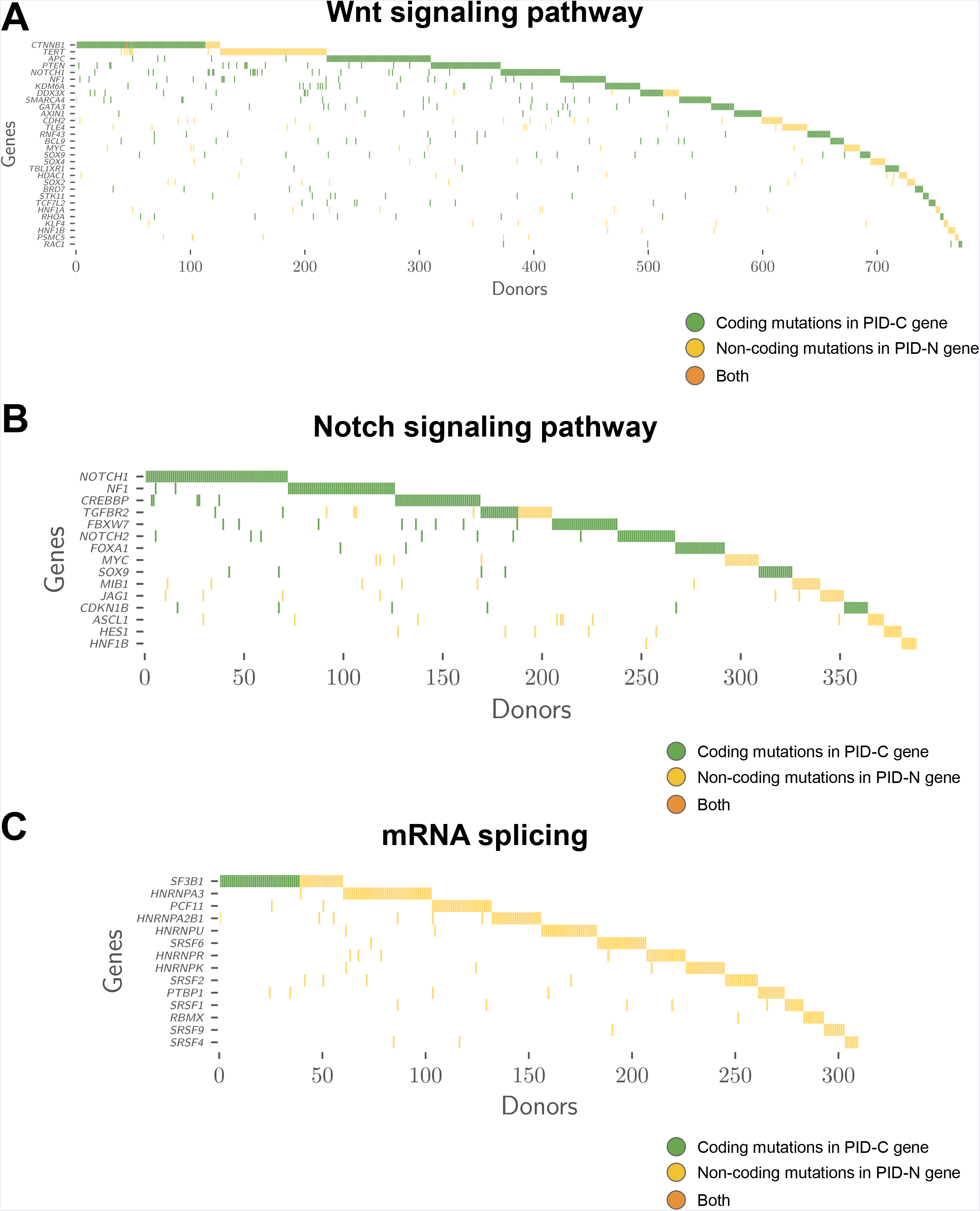
Oncoprints for mutations in biological pathways and processes identified by pathway and network methods. **(A) Oncoprint for Wnt signaling pathway altered by both coding and non-coding mutation**. Coding mutations in PID-C genes in the Wnt signaling pathway (GO:0016055) occur in 606 tumors, and non-coding mutations in PID-N genes in the Wnt signaling cover an additional 169 tumors (additional 28% tumors). **(B) Oncoprint for Notch signaling pathways altered by both coding and non-coding mutations**. Coding mutations in PID-C genes in the Notch signaling pathway (GO:0007219) occur in 304 tumors, and non-coding mutations in PID-N in the Notch signaling pathway genes cover an additional 85 tumors (additional 29% tumors). **(C) Oncoprints for pathways enriched by non-coding mutations in RNA Splicing**. Coding mutations in the PID-C gene *SF3B1* in the “mRNA splicing via splicesome” (GO:0000398) pathway occur in 39 tumors, while non-coding mutations in 15 PID-N genes in the same pathway cover an additional 271 tumors.

**Figure S13:**
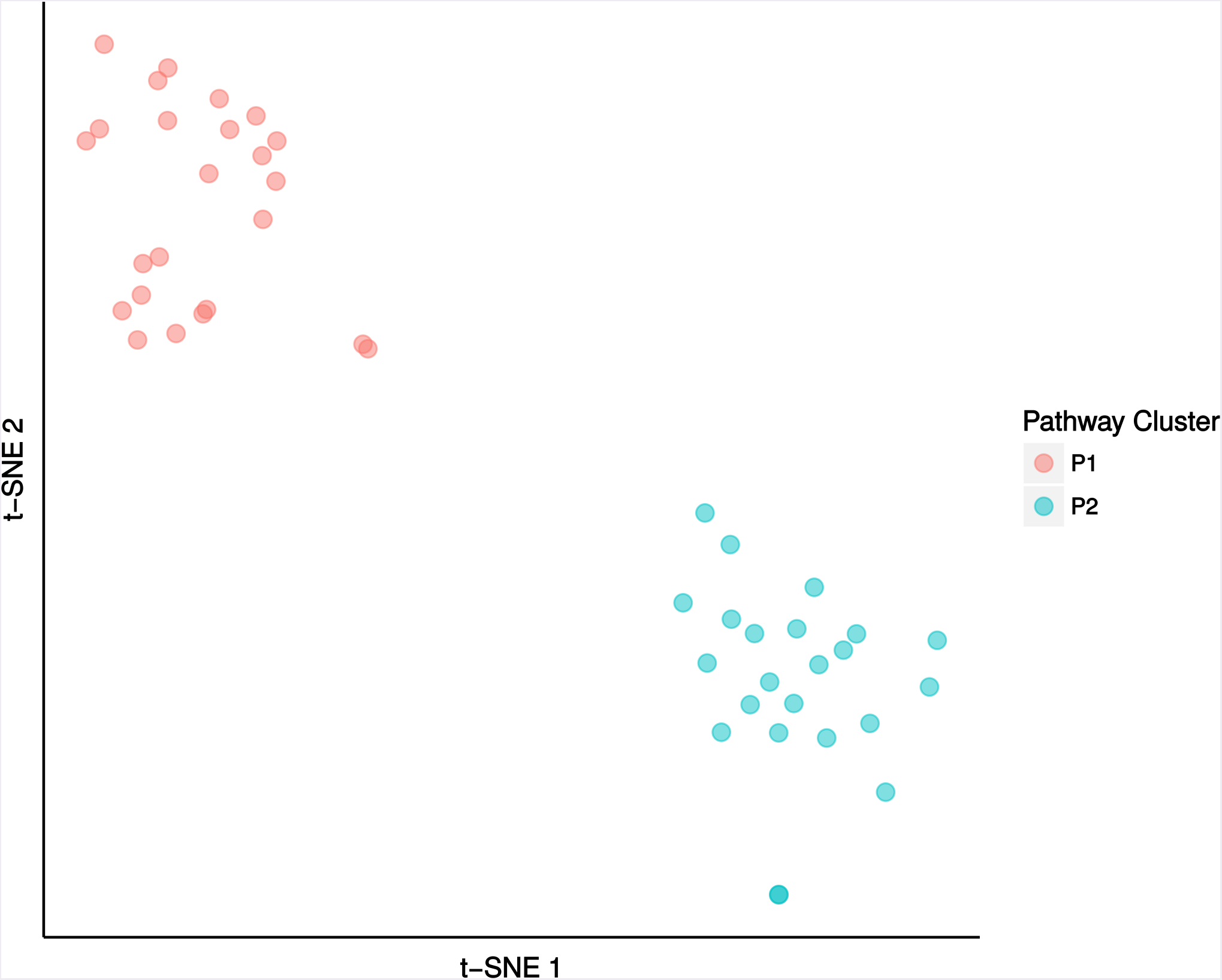
tSNE plot of pathway enrichment scores. Clustering of 47 curated pathway^32^ into two distinct pathway clusters.

## Online Methods

### Mutation and Pathway Data

We combined several pathways and interaction networks with gene scores derived from the PCAWG drivers analysis^1^ for use by pathway and networks methods. Here, we use the term “pathway methods” for those approaches that make use of sets of related genes for their analysis while the term “network methods” are reserved for those that also incorporate the interactions among the genes and/or their products.

#### Somatic mutation data

We obtained consensus driver *p*-values (syn8494939) from the PCAWG drivers analysis^1^ for coding and non-coding (core promoter, 5’ UTR, 3’ UTR, enhancers) genomic elements for the Pancan-no-skin-melanoma-lymph cohort. We removed driver *p*-values for several elements (*H3F3A* and *HIST1H4D* coding; *LEPROTL1, TBC1D12*, and *WDR74* 5’ UTR; and chr6:142705600-142706400 enhancer, which targets *ADGRG6*) that the PCAWG drivers analysis had manually examined and discarded. We included enhancers with ≤ 5 gene targets (syn7188184), which covered 89% of enhancers elements from the PCAWG drivers analysis^1^. In cases where the PCAWG drivers analysis reported multiple *p*-values for the same genomic element, we used the smallest reported *p*-value for that element.

#### Derivation of gene scores

Pathway databases and gene interaction networks typically record information at the level of individual genes. Thus, we formed coding and non-coding gene scores by combining PCAWG driver *p*-values across coding and/or non-coding (core promoter, 5’ UTR, 3’ UTR, enhancer) genomic elements as follows. Let *p_x_*(*g*) be the driver *p*-value for element x of gene *g* from the PCAWG drivers analysis^1^. We combined *p*-values from multiple elements using Fisher’s method, where we selected the minimum *p*-value min(*p_promoter_*(*g*), *p*_5’*UTR*_(*g*)) for overlapping core promoter and 5’ UTR elements on gene *g* and the minimum *p*-value *p_enhancer_*(*g*) of all enhancers targeting gene g. Using this approach, we defined the following gene scores on coding (GS-C), non-coding, (GS-N), and combined coding and non-coding (GS-CN) genomic elements:

1. GS-C: *p_c_*(*g*) = *p_coding_*(*g*)
2. GS-N: *p*_N_(*g*) = fisher(min(*P*_promoter_(*g*), *P*_5’UTR_(*g*)), *P*_3’UTR_(*g*), *P*_enhancer_(*g*))
3. GS-CN: *p*_CN_(*g*) = fisher(*P*_coding_(*g*), min(*P*_promoter_(*g*), *P*_5’UTR_(*g*)), *P*_3’UTR_(g), *P* enhancer^(g))^

Here, *p* = fisherp, …, *p_k_*) is Fisher’s method, 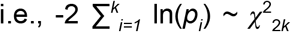, for independently and identically distributed *p*_1_, …, *p_k_* ~ *U*(0, 1), where *2k* is the degrees of freedom in the calculation. Moreover, when the driver *p*-value for a genomic element was undefined, we omitted that element from the calculation and reduced the number of degrees of freedom.

For the pathway and networks methods that analyze individual mutations, we used mutations from the PCAWG MAF (syn7118450) on the same genomic elements (syn5259890) as the PCAWG drivers analysis, i.e., coding, core promoter, 5’ UTR, 3’ UTR, and enhancer. We removed melanoma and lymphoma samples as well as 69 hypermutated samples with over 30 mutations/MB (syn7222520, syn7814911). We also removed mutations in elements that the PCAWG drivers analysis had manually examined and discarded (see above), resulting in lists of mutations used for later assessing biological relevance of our results (syn8103141, syn9684700).

#### Pathway and network databases

Pathway methods used gene sets from six databases: CORUM^2^ (syn11426307), GO^3,4^ (syn3164548), InterPro^5^ (syn11426307), KEGG^6^ (syn11426307), NCI Nature^7^ (syn11426307), and Reactome^8^ (syn3164548), where small (< 3 genes) and large (> 1,000 genes) pathways were removed.

Network methods used interactions from three interaction networks: the largest connected subnetwork of the ReactomeFI 2015 interaction network^9^ (syn3254781) with high-confidence (≥ 0.75 confidence score) interactions, which we treated as undirected; the largest connected subnetwork of the iRefIndex14 interaction network^10^, which we augmented with interactions from the KEGG pathway database^6^ (syn10903761); and the largest connected subnetwork of the STRING v10 network^11^ (syn11712027) with high-confidence (> 9 confidence score) interactions. The BioGRID interaction network^12^ (syn3164609) was also used to evaluate and annotate results.

### Pathway and Network Integration of Gene-Level Scores

#### Individual pathway and network algorithms

We applied seven pathway and network methods to the gene scores and mutation data. We used two pathway methods: ActivePathways [Paczkowska, Barenboim, *et al*., in submission] and a hypergeometric analysis [Vazquez]. We also used five network methods: CanIsoNet [Kahraman *et al*., in preparation], Hierarchical HotNet^13^, an induced subnetwork analysis [Reyna and Raphael, in preparation], NBDI^14^, and SSA-ME^15^.

**Table M1:**
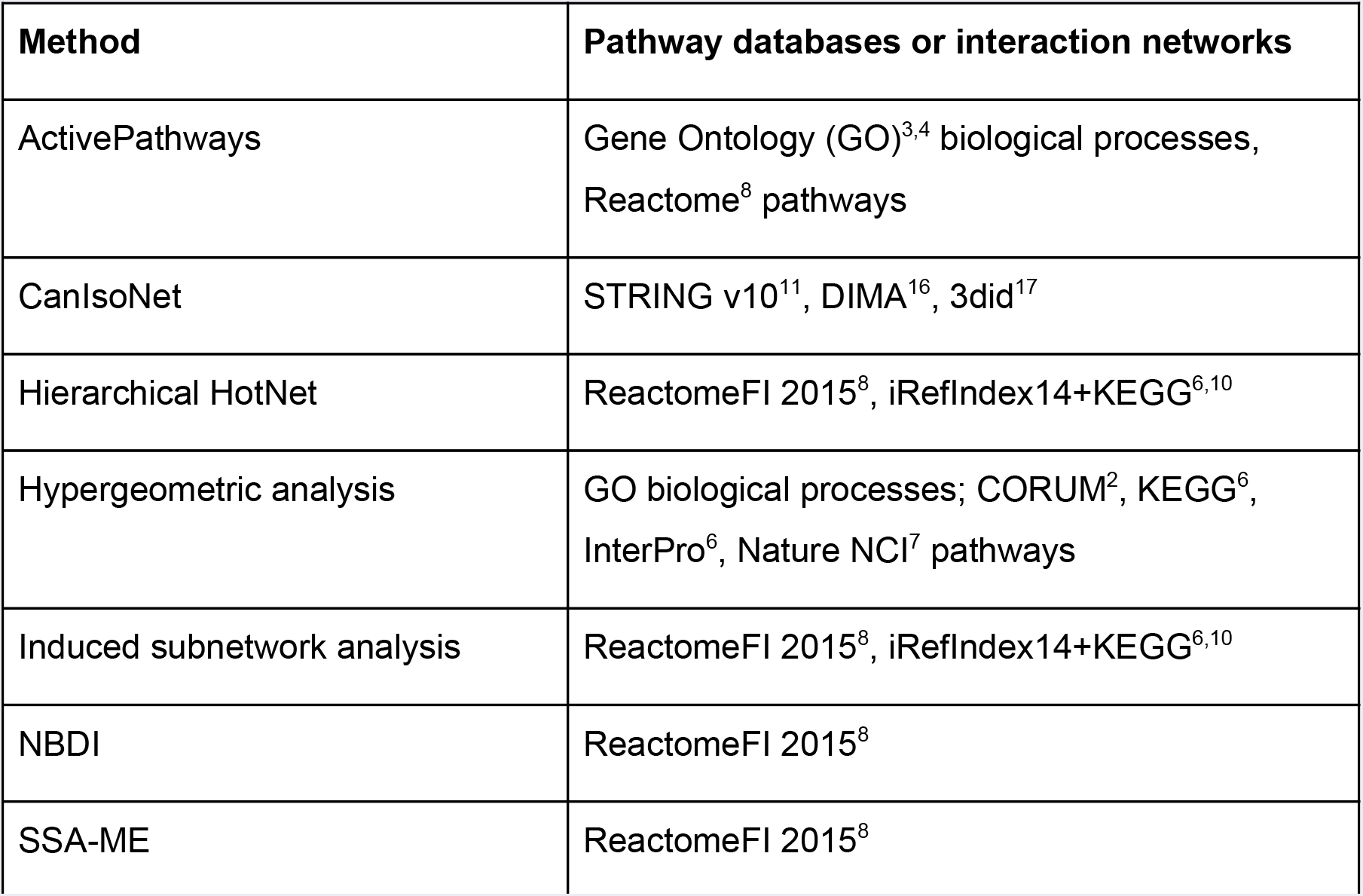
Summary of pathway database and interaction network data for each method.

| Method | Pathway databases or interaction networks |
| --- | --- |
| ActivePathways | Gene Ontology (GO)<sup>3,4</sup> biological processes, Reactome<sup>8</sup> pathways |
| CanIsoNet | STRING v10<sup>11</sup>, DIMA<sup>16</sup>, 3did<sup>17</sup> |
| Hierarchical HotNet | ReactomeFI 2015<sup>8</sup>, iRefIndex14+KEGG<sup>6,10</sup> |
| Hypergeometric analysis | GO biological processes; CORUM<sup>2</sup>, KEGG<sup>6</sup>, InterPro<sup>6</sup>, Nature NCI<sup>7</sup> pathways |
| Induced subnetwork analysis | ReactomeFI 2015<sup>8</sup>, iRefIndex14+KEGG<sup>6,10</sup> |
| NBDI | ReactomeFI 2015<sup>8</sup> |
| SSA-ME | ReactomeFI 2015<sup>8</sup> |

Using these pathway and network databases, we ran each method on the GS-C, GS-N, and GS-CN gene scores to identify three corresponding lists of genes. Each method evaluated the statistical significance of its results on each dataset.

#### Non-coding value-added (NCVA) procedure

The GS-CN results leverage both coding and non-coding mutation data, improving the detection of weaker pathway and network signals. We devised a non-coding value-added (NCVA) procedure to separate the coding and non-coding signals in this combined analysis, resulting in a set of NCVA genes for which the non-coding mutation data makes a statistically significant contribution to their discovery in the GS-CN results. Specifically, we evaluated the statistical significance of genes in the GS-CN results using a permutation test where the driver *p*-values for coding elements were fixed and the driver *p*-values for non-coding elements were permuted. This procedure identified the subset of the GS-CN results that were reported infrequently (*p* < 0.1) on permuted data and thus more likely to be true positives. Each method’s NCVA results were added to that method’s overall set of non-coding results (GS-N).

#### Consensus results for pathway and network methods

We defined a consensus set of genes for each set of results: GS-C results, GS-N results, GS-CN results, and GS-N combined with NCVA results, across our seven pathway and network methods. Specifically, we defined a gene to be a consensus gene if it was found by a majority (≥ 4/7) of the pathway and network methods. For our analysis, we focused on the consensus GS-C results, which we call the pathway-implicated driver genes with coding variants (PID-C), and the consensus from the GS-N results combined with NCVA results, which we call the pathway-implicated driver genes with non-coding variants (PID-N). We defined PID-C genes as the 87 genes in the consensus of the GS-C results, and we defined PID-N genes as the 93 genes in the consensus of each method’s GS-N results combined with its NCVA results.

### Downstream Interpretation of Pathway-Implicated Drivers

We performed several analyses to assess the biological relevance of PID-C and PID-N genes.

#### Identification of mutational signatures of PID genes

We performed a permutation-based enrichment test for mutation signatures from PCAWG mutation signatures analysis^18^. We identified the most likely mutation signature for each non-coding mutation in PID-N genes and compared them to randomly chosen non-coding mutations in non-PID-N genes.

#### Gene scores improve network neighborhood scores of PID genes

To assess the extent to which gene scores on PID genes contribute to their detection by pathway and network methods, we considered the contribution of each PID gene’s score to the score of its network neighborhood in the BioGRID interaction network.

For each PID gene g, we used Fisher’s method to combine the gene scores of the first-order network neighbors of *g* both with and without the score of *g* itself. In particular, for gene *g*, let *p*(*g*) be the gene score for *g* and *N*(*g*) be the network neighborhood of g. Then

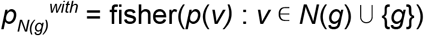

is a score for the network neighborhood of *g* when *including* gene *g* and

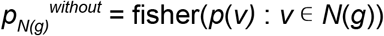

is a score for the network neighborhood of *g* when *excluding* gene *g*.

If the network neighborhood of *g* has a smaller *p*-value with *g* than without *g*, i.e., *p_N(g)_^wlth^* > *p_N(g)_^wlthout^*, then gene *g improves* the score of the network neighborhood, suggesting that the gene score of *g* plays a role in its detection by pathway and network methods. Alternatively, if the network neighborhood of *g* has a larger *p*-value with *g* than without *g*, i.e., *p_N(g)_^with^* > *p*_N(g)_^without^, then gene *g worsens* the score of the network neighborhood, suggesting that the gene scores of the network neighbors of *g* are predominantly responsible for the detection of *g* by pathway and network methods.

We performed this test for every PID-C gene with GS-C gene scores and every PID-N gene with GS-N gene scores. We also sampled genes uniformly at random from the network (87 for PID-C genes and 93 for PID-N genes; 10^6^ trials) to ascertain whether significantly more PID genes that improved the scores of their network neighborhoods than expected by chance.

#### Expression analysis of PID genes

We evaluated whether mutation status of each PID gene was correlated with RNA expression. We used PCAWG-3 gene expression data (syn5553991), which was averaged from TopHat2 and STAR-based alignments, with FPKM-UQ normalization. Tumor type and copy-number aberrations are known to be covariates for gene expression, so we conditioned on tumor types and annotated copy-number aberrations.

We used the following procedure to assess expression correlations on individual tumor types. We only considered cases with at least 3 mutated samples and 3 non-mutated samples to restrict our analysis to cases with sufficient statistical power. For each PID-C gene or each non-coding element in a PID-N gene, we partitioned the samples with expression data into a set *A* of samples with mutation(s) in the element and a set *B* of samples without mutations in the element. We performed the Wilcoxon rank-sum test for the expression of the gene in sets *A* and *B* and performed the Benjamini-Hochberg correction on each coding or non-coding element to provide FDRs.

We used the following procedure to assess expression correlations across tumor types. We only considered cases with at least 1 mutated sample and 1 non-mutated sample to restrict our analysis to cases with sufficient statistical power. For each PID-C gene and each non-coding element in a PID-N gene, we partitioned the samples with expression data into sets *A_c_* of samples in cohort *c* with mutation(s) in the element and sets *B_c_* of samples in cohort *c* without mutations in the element. We converted the expression values into z-scores using the expression from non-mutated samples in cohort c, and we computed the Wilcoxon rank-sum test on the expression of the gene in sets from *A* = ⋃_*c* ∊ *C*_ *A_c_* and *B* = ⋃_*c* ∊ *C*_ *B_c_*, where *C* is the set of all cohorts containing samples with mutation(s) in the element. We then performed the Benjamini-Hochberg correction on each coding or non-coding element to provide FDRs.

#### Network annotation of PID genes

We performed a permutation test to evaluate the statistical significance of the number of interactions in the BioGRID high-confidence interaction network between PID-C genes, the number of interactions between PID-N genes, and the number of interactions between PID-C and PID-N genes, i.e., when a PID-C gene interacts with a PID-N gene. To compute the permutation *p*-value we sampled random networks uniformly at random from the collection of networks with the same degree sequence as the BioGRID network.

We found connected subnetworks of 46 PID-C genes (31 genes expected, *p* = 9 × 10^-4^) and 16 PID-N genes (10 genes expected, *p* = 6.1 × 10^-2^) in the high-confidence BioGRID^19^ protein-protein interaction (PPI) network. The union of the PID-C and PID-N genes formed a larger connected subnetwork of 73 genes (**Figure 4A**). These connected subnetworks were significantly larger than expected by chance according to this permutation test (57 genes expected, *p* = 2.2 × 10^-3^). Further, we observed statistically significant numbers of protein-protein interactions between PID-C and PID-N genes (67 interactions observed vs. 45 expected, *p* = 6 × 10^-4^), suggesting that the associated mutations may target an overlapping set of underlying pathways. The PID-C genes were connected by significantly more interactions than expected (64 vs. 40 expected, *p* < 10^-4^) and the PID-N genes were interconnected at a sub-significant level (18 vs 12 expected, *p* = 6.8 × 10^-2^). Thus certain pathways are affected by either coding or non-coding mutations, but some pathways are affected by a complement of both coding and non-coding mutations.

#### Pathway annotation of PID genes

Using g:Profiler^20^, we performed a pathway enrichment analysis for PID genes and 12,061 gene sets representing GO biological processes and Reactome pathways. We used the Benjamini-Hochberg correction to control the FDR of the results.

#### Characterization of PID genes in RNA splicing

GSEA enrichment analysis was performed with the default parameters using the curated pathway gene lists^21^ for samples harboring non-synonymous coding mutations in 5 genes (*FUBP1, RBM10, SF3B1, SRSF2*, and *U2AF1*) with confirmed on-target splicing deregulation. Due to limited number of samples with RNA-seq data in individual tumor types, we restricted our analysis to missense mutations in *SF3B1*, truncating mutations in *RBM10*, and truncating mutations in *FUBP1* for tumor types contained at least 3 samples with these classes of mutations. Each tumor type containing such mutations was considered separately^21^.

We performed the same GSEA analysis for non-coding mutations in 17 PID-N genes that were annotated as involved in RNA splicing. Due to limited number of samples from individual tumor types containing mutations in these genes (often there was only 1 per tumor type), we performed GSEA analysis jointly on all tumor types containing mutations in an individual PID-N gene, restricting the non-mutated group to samples from the same tumor types as the mutant samples. The GSEA Normalized Enrichment Scores (NES) were clustered using hierarchical complete linkage clustering on the Euclidean distance between the NES scores. Separately, we computed a 2D projection of NES scores using t-Distributed Stochastic Neighbor Embedding (tSNE).

### Additional Information

See **Supplement** for more information about data processing and details of individual network and pathway methods.

